# A learning-evoked slow-oscillatory architecture paces population activity for offline reactivation across the human medial temporal lobe

**DOI:** 10.64898/2026.02.12.705512

**Authors:** Adrien A. Causse, Jonathan Curot, Vítor Lopes-dos-Santos, Raphaël Nunes-da-Silva, Helen C. Barron, Vincent Dornier, Marie Denuelle, Amaury De Barros, Jean-Christophe Sol, Jean-Albert Lotterie, Katia Lehongre, Sara Fernandez-Vidal, Valerio Frazzini, Vincent Navarro, Luc Valton, Emmanuel J. Barbeau, Tim Denison, Leila Reddy, David Dupret

## Abstract

Memory processing requires the coordinated engagement of neuronal populations across distributed brain networks and across time. How such coordination is organized in the human medial temporal lobe (MTL) remains unclear. Here, we show that MTL population activity is dynamically structured by a transient slow-oscillatory architecture that emerges during online learning to promote offline consolidation and later recall. Using intracranial recordings that combine single-neuron spiking activity and local field potentials in human participants, we find that mnemonic engagement evokes on-demand slow-oscillatory bursts in the hippocampus. These hippocampal bursts synchronize gamma-band patterns across MTL regions, defining discrete coordination events that pace cross-regional coactivity motifs during learning. These learning-time population motifs are then selectively reactivated during hippocampal ripples in post-learning rest, and the strength of their reactivation predicts subsequent recall accuracy. Together, these findings identify a multi-scale coordination mechanism that links distributed population activity across learning, consolidation, and recall in humans.

## Introduction

Memory unfolds across an extended processing arc that spans experience and time, from learning through consolidation to recall ^1–3^. At the core of the brain memory circuitry is the hippocampus, whose neurons support the representation of relationships among stimuli and events ^4–8^. During learning, hippocampal activity gives rise to structured patterns of neuronal coactivity that are subsequently reactivated offline during rest and later reinstated online during recall ^9–11^.

Although these processes have been extensively characterized at the behavioral and representational levels, the population-level network mechanisms that dynamically coordinate hippocampal activity across learning, consolidation, and recall in humans remain unclear.

Coordinating memory across this processing arc requires interactions among the hippocampus and the broader medial temporal lobe (MTL) network ^12,13^. The hippocampus interacts with entorhinal, parahippocampal, and amygdala regions to support memory.

Integrating activity across this multi-region system poses a fundamental challenge. Memory-related processing requires integration of neuronal spiking, population-level synchronization, and interregional communication across multiple temporal and spatial scales, while preserving local computational dynamics within individual regions. Achieving such coordination demands mechanisms capable of transiently binding distributed neural activity into coherent functional states and organizing population-level communication across regions. Oscillatory activity provides a powerful means of achieving this coordination by defining temporal windows in which patterns of neuronal coactivity form, interact, and become eligible for subsequent offline consolidation and later reinstatement.

In animal models, hippocampal oscillations have been shown to provide a temporal framework for coordinating memory-related population activity^14^. In rodents, theta-band (5–12 Hz) oscillations are readily observed as rhythmic fluctuations in local field potentials (LFPs) during voluntary movement and spatial exploration^15,16^. These oscillations provide a temporal scaffold for organizing millisecond-timescale patterns of neuronal coactivity during learning and their reinstatement during memory-guided behavior^14,17,18^. Theta-nested gamma-band oscillations (30–150 Hz) have been proposed to reflect both local population computation within the hippocampus and communication channels through synchronization with distributed brain regions^19–21^. Patterns of neuronal coactivity formed during theta-governed online states are subsequently reactivated during high-frequency ripple events (100–250 Hz) that occur during rest and sleep, supporting offline memory consolidation^9,17,18,22–25^.

Together, these findings have shaped influential models of memory and cognition in which oscillatory dynamics link online processing during awake behavior with subsequent consolidation during offline states^7,10,26–29^. However, accumulating evidence indicates that hippocampal oscillatory dynamics vary substantially across species and behavioral contexts. In larger-brained mammals, including rabbits, cats, bats, and primates, hippocampal rhythmic activity is often slower, more intermittent, and more closely linked to task demands than to behavioral exploration ^30–33^. In humans, intracranial recordings frequently indicate lower-frequency activity whose functional significance remains elusive^34–38^. As a result, it is unclear whether the human MTL expresses a unifying coordination architecture capable of linking neuronal spiking, network synchronization, and offline reactivation across memory processing stages.

Here we show that human memory is organized by a coordination architecture in which brief, task-evoked rhythmic events link neuronal spiking, interregional synchronization, and offline reactivation across the full memory processing arc. Using intracranial electroencephalography combined with simultaneous single-neuron recordings from the human MTL during a relational memory task, we identify a coordination architecture that emerges as transient, task-evoked 2-Hz oscillatory bursts during learning and recall. These oscillatory events pace neuronal population spiking and synchronize gamma-band activity across hippocampal and extra-hippocampal MTL regions, defining temporal windows for coordinated network interactions. Critically, coactivity motifs structured by this oscillatory architecture during learning are selectively reactivated during hippocampal ripple events in post-learning rest. The strength of this reactivation predicts subsequent recall accuracy, directly linking oscillatory coordination during learning to offline consolidation and later memory performance. Together, these findings reveal a coordination architecture that operates through discrete, mnemonic engagement-locked events rather than as a sustained background rhythm, coordinating MTL activity across the full arc of memory processing, from learning through consolidation to recall in humans.

## Results

### Mnemonic engagement evokes transient slow oscillatory activity in the human hippocampus

To examine how hippocampal network dynamics evolve across the full memory processing arc in humans, we designed an associative relational task that enabled continuous intracranial recordings from learning through consolidation to recall (Figure 1A). In this task, we trained 27 participants to learn associations among individuals in a community. Each recording day began with a pre-learning rest (pre-rest), followed by a viewing session during which participants were familiarized with photographs of community members presented in random order. During the subsequent learning session, participants learned associations between pairs of individuals (for example, “*Marie knows Antoine*”). Learning progress was assessed through intermittent multiple-choice questions confirming knowledge of the community structure (Figure 1B). After a post-learning rest (post-rest), participants completed a memory recall test session that confirmed successful retention of the learned associations (Figure 1C; *P* < 0.001; Wilcoxon signed-rank test compared to 33% chance level).

**Figure 1.**
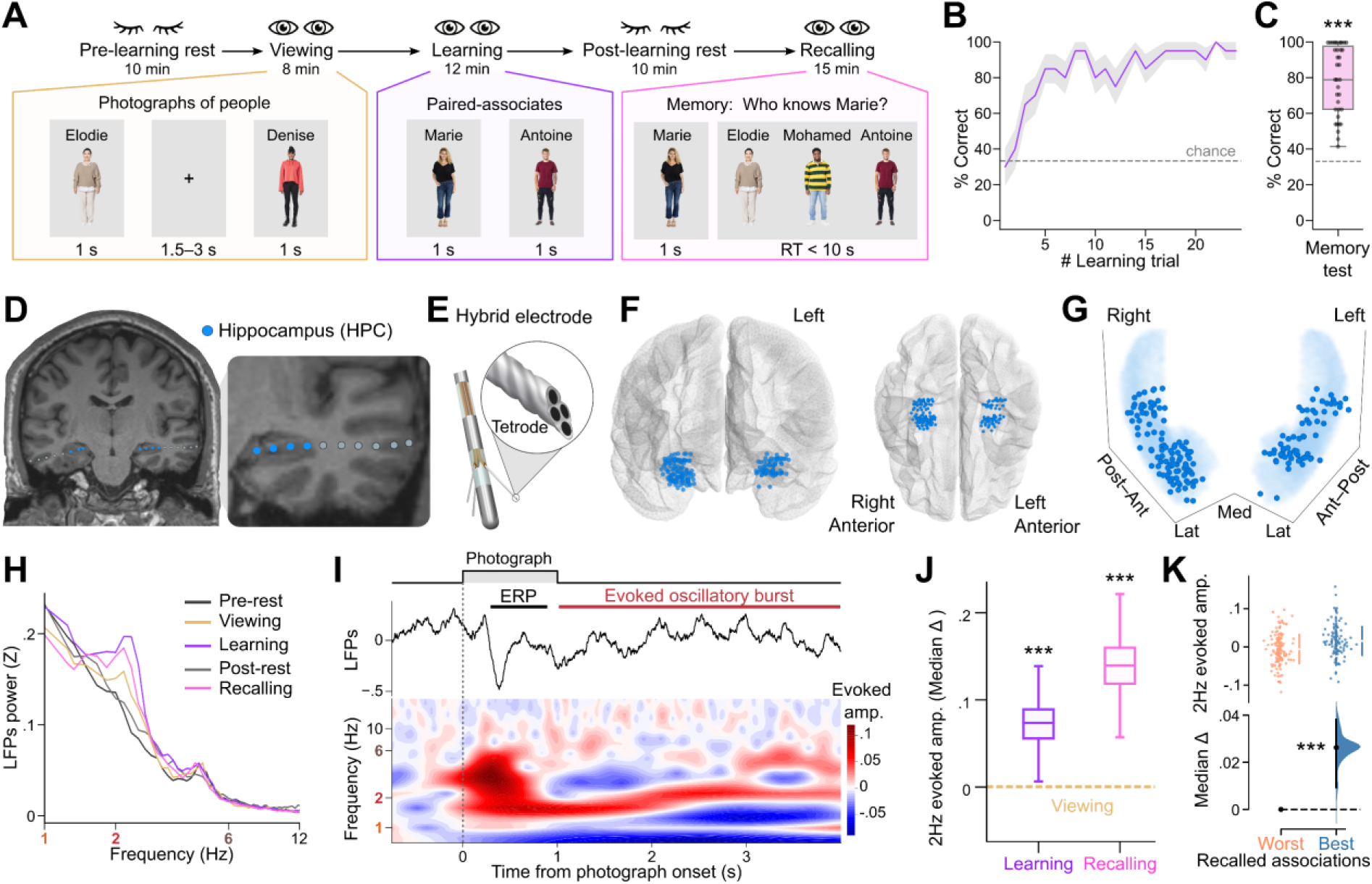
Mnemonic engagement evokes transient slow oscillatory activity in the human hippocampus **(A)** Schematic of the relational memory task with example stimulus photographs. **(B)** Learning performance (percentage correct trials). **(C)** Memory recall accuracy (percentage correct responses). **(D)** T1-weighted MRI showing contact locations from two representative depth electrodes targeting the hippocampus. **(E)** Hybrid micro-/macro-electrode design with tetrodes extending from the macrocontact shaft. **(F,G)** MNI brain template **(F)** and 3D projection **(G)** of hippocampal contacts across participants (axes: Post–Ant, posterior–anterior; Med–Lat, medio–lateral; Sup–Inf, superior–inferior). **(H)** Power spectral density (PSD) from an example hippocampal contact across task stages, illustrating task-dependent modulation of slow oscillatory activity. **(I)** Peri-stimulus average of hippocampal LFPs (top) and corresponding spectrogram (bottom), showing a stimulus-locked event-related potential (ERP) followed by transient slow oscillatory activity during the inter-stimulus interval. **(J)** Median difference in evoked 2-Hz oscillatory amplitude during learning and recall relative to viewing, computed over post-ERP epochs (>1 s after photograph onset). **(K)** Estimation plot showing the difference in amplitude of evoked 2-Hz oscillatory bursts during learning trials associated with best versus worst subsequent memory recall. Upper panel: raw data (points) with mean ± SD (vertical lines); bottom panel: mean difference (black dot) with 95% confidence interval (black ticks) and bootstrapped sampling-error distribution (filled curve) relative to the worst-recalled associations reference (horizontal dashed line). Data were analyzed using two-sided paired permutation tests, except in **C**, where a Wilcoxon signed-rank test was applied; ***P < 0.001.

Intracranial recordings were obtained using hybrid depth electrodes that enabled simultaneous measurement of hippocampal LFPs and population-level single-neuron spiking (Figures 1D–G and S1A-C). Each electrode shaft incorporated multichannel tetrodes (Figure 1E), similar to those commonly employed in rodent studies ^39,40^, combined with macrocontacts for clinical epilepsy monitoring ^41^. To evaluate the network expression of a coordination architecture able to link hippocampal activity across memory stages, we decomposed LFP signals into their constituent oscillatory components using an unsupervised, data-driven approach that does not impose predefined frequency bands, previously validated in rodents (Figure S1C-F) ^39,40^. Using this approach, we identified a prominent slow oscillatory component centered near 2 Hz [peak (80% power band): 2.38 (1.25–3.50) Hz], alongside a slower (∼1 Hz) and a faster (∼6 Hz) rhythmic component (Figure S1C–F). Notably, standard local (bipolar) referencing markedly reduced sensitivity to this slow oscillatory component (Figure S1G-I), and its expression was strongest at electrode contacts outside interictal zones (Figure S1J,K)^42^, providing a potential explanation for why it has been difficult to detect in prior human studies.

Mnemonic engagement during learning and recall was associated with enhanced expression of this slow oscillatory component in the hippocampus (Figure 1H,I). Hippocampal 2-Hz power was significantly greater during both learning and recall than during resting or viewing sessions (learning, *P* < 0.001; recalling, *P* = 0.006; paired permutation tests compared to pre-learning rest), with no comparable modulation observed at adjacent slower or faster frequencies (Figure S1L,M). Linear mixed-effects modeling confirmed that this enhancement was selective for the 2-Hz oscillatory component (Figure S1N).

This slow oscillatory activity was expressed as brief, stimulus-locked bursts (mean burst duration [95% confidence interval (CI)]: 19.6 [15.8 – 23.4] cycles per burst; Figure S1O-Q), rather than as sustained rhythmic activity. Peri-stimulus averages of hippocampal LFPs revealed a photographic stimulus-locked event-related potential (ERP), with transient slow oscillatory activity emerging during the inter-stimulus interval (Figures 1I and S1R). The amplitude of this slow oscillatory burst activity was significantly enhanced during learning and recall (Figures 1J and S1S; learning, P < 0.001; recalling, P < 0.001; paired permutation tests compared to viewing), and correlated with the magnitude of the ERP deflection (Figure S1T), with no comparable modulation for neighboring 1-Hz or 6-Hz bands (Figure S2T,U). Critically, photographs that elicited the strongest slow oscillatory responses during learning were those subsequently best remembered (Figure 1K; best versus worst recalled associations, P < 0.001; paired permutation tests).

These findings show that mnemonic processing selectively evokes in the human hippocampus a slow oscillatory response that emerges as transient, stimulus-locked activity, and that the magnitude of this response predicts subsequent memory performance.

### Learning-evoked slow-oscillatory architecture paces hippocampal population spiking and gamma-band activity

To characterize the neural architecture underlying memory-evoked slow oscillatory activity, we examined how hippocampal neuronal spiking and local network dynamics relate to the phase of this oscillatory signal (Figure 2A). At the population level, hippocampal spiking activity exhibited clear modulation by slow oscillatory phase. Average hippocampal LFPs aligned to oscillatory phase revealed rhythmic modulation of population firing rates at the slow oscillatory timescale (Figures 2B and S2A-C). Across neurons, spikes were more strongly phase-locked to the slow oscillatory rhythm (2-Hz) than to slower (1-Hz) or faster (6-Hz) comparison frequencies (Figure 2C,D; 2-Hz versus 1-Hz, *P* = 0.003; 2-Hz versus 6-Hz, *P* = 0.020; 1-Hz versus 6-Hz, *P* = 0.815; two-sided paired permutation tests), indicating selective coordination of spiking by the slow oscillatory signal.

**Figure 2.**
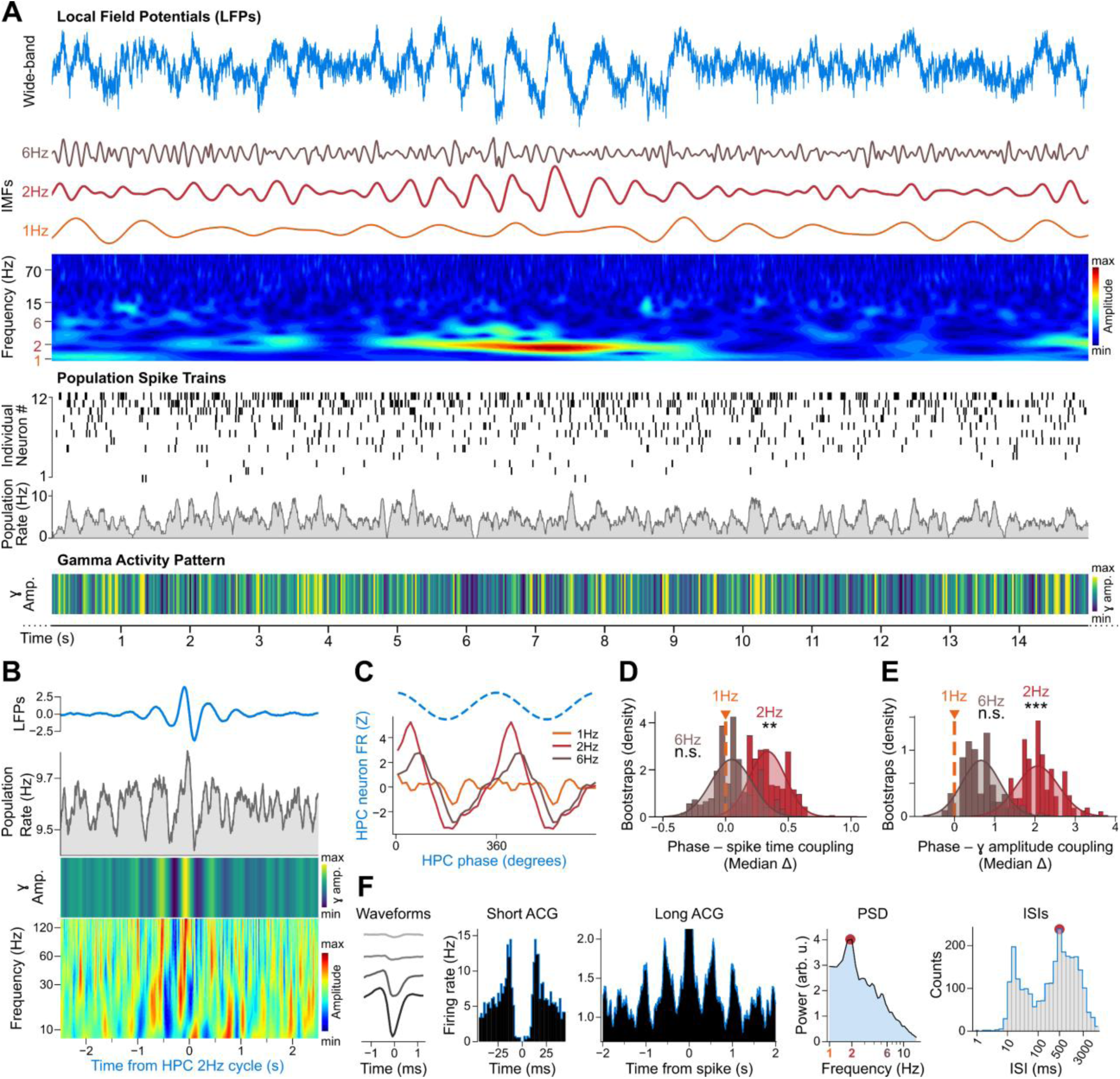
A slow oscillatory architecture paces hippocampal spiking and gamma activity **(A)** Example hippocampal tetrode recording illustrating slow oscillatory activity, neuronal spiking, and gamma-band activity. From top to bottom: wideband LFP trace decomposed into its constituent oscillatory components (intrinsic mode functions, IMFs) with spectrogram; raster plot of single-neuron spike trains with population firing rate; and corresponding gamma-band activity pattern. **(B)** Average hippocampal LFPs aligned to 2-Hz oscillatory phase with instantaneous population firing rate, gamma amplitude, and spectrogram, showing rhythmic modulation. **(C)** Firing-phase histograms of an example neuron relative to 1-, 2-, and 6-Hz oscillations. **(D)** Spike–phase consistency differences between 1-Hz and 2-or 6-Hz oscillations across hippocampal neurons. **(E)** Phase–amplitude coupling differences for hippocampal gamma envelopes between 1-Hz and 2-or 6-Hz oscillations. **(F)** Representative hippocampal neuron exhibiting slow oscillatory rhythmicity. From left to right: mean spike waveform across tetrode channels, short-and long-timescale spike autocorrelograms (ACGs), power spectral density (PSD), and inter-spike interval (ISI) distribution. Statistics were assessed using two-sided paired permutation tests; ***P < 0.001, **P < 0.01; n.s., not significant.

Gamma-frequency activity is thought to reflect local population spiking subspaces in animal models^21^. Consistent with this, human hippocampal spiking correlated more strongly with local than with distal gamma-band (60–160 Hz) activity (Figure S2D). The amplitude of hippocampal gamma oscillations was more strongly modulated by slow oscillatory phase than by the phase of slower (1-Hz) or faster (6-Hz) rhythms (Figure 2B,E; 2-Hz versus 1-Hz, *P* < 0.001; 2-Hz versus 6-Hz, *P* < 0.001; 1-Hz versus 6-Hz, *P* = 0.097; two-sided paired permutation tests), indicating selective cross-frequency coupling. This modulation extended along the hippocampal anteroposterior axis (Figure S2E-K).

At the single-neuron level, hippocampal neurons also exhibited prominent slow-timescale rhythmicity in their spike train autocorrelograms (Figures 2F and S2L,M), confirming that slow oscillatory structure was expressed at both population and single-neuron scales.

Together, these findings show that memory-evoked slow oscillatory activity in the human hippocampus constitutes a coherent temporal architecture that paces population spiking and organizes local gamma-band network dynamics.

### The hippocampal slow-oscillatory architecture synchronizes activity across the MTL

We next asked whether this hippocampal architecture extends beyond the hippocampus to coordinate activity across the MTL network. To address this, we examined intracranial recordings from electrodes targeting other MTL regions (for example, entorhinal cortex) as well as regions outside the MTL (for example, temporal cortices) (Figure 3A,B). As in the hippocampus (Figure 2F), single-neuron spike trains recorded across MTL regions exhibited slow oscillatory rhythmicity centered near 2 Hz (Figure 3C), whereas neurons recorded outside the MTL displayed rhythmicity at other frequencies (Figure 3C). Spiking activity in MTL neurons was more strongly phase-locked to hippocampal slow oscillatory phase (2-Hz) than to slower or faster rhythms (Figure 3D,E; 2-Hz versus 1-Hz, *P* = 0.002; 2-Hz versus 6-Hz, *P* < 0.001; 1-Hz versus 6-Hz, *P* = 0.059; two-sided paired permutation tests).

**Figure 3.**
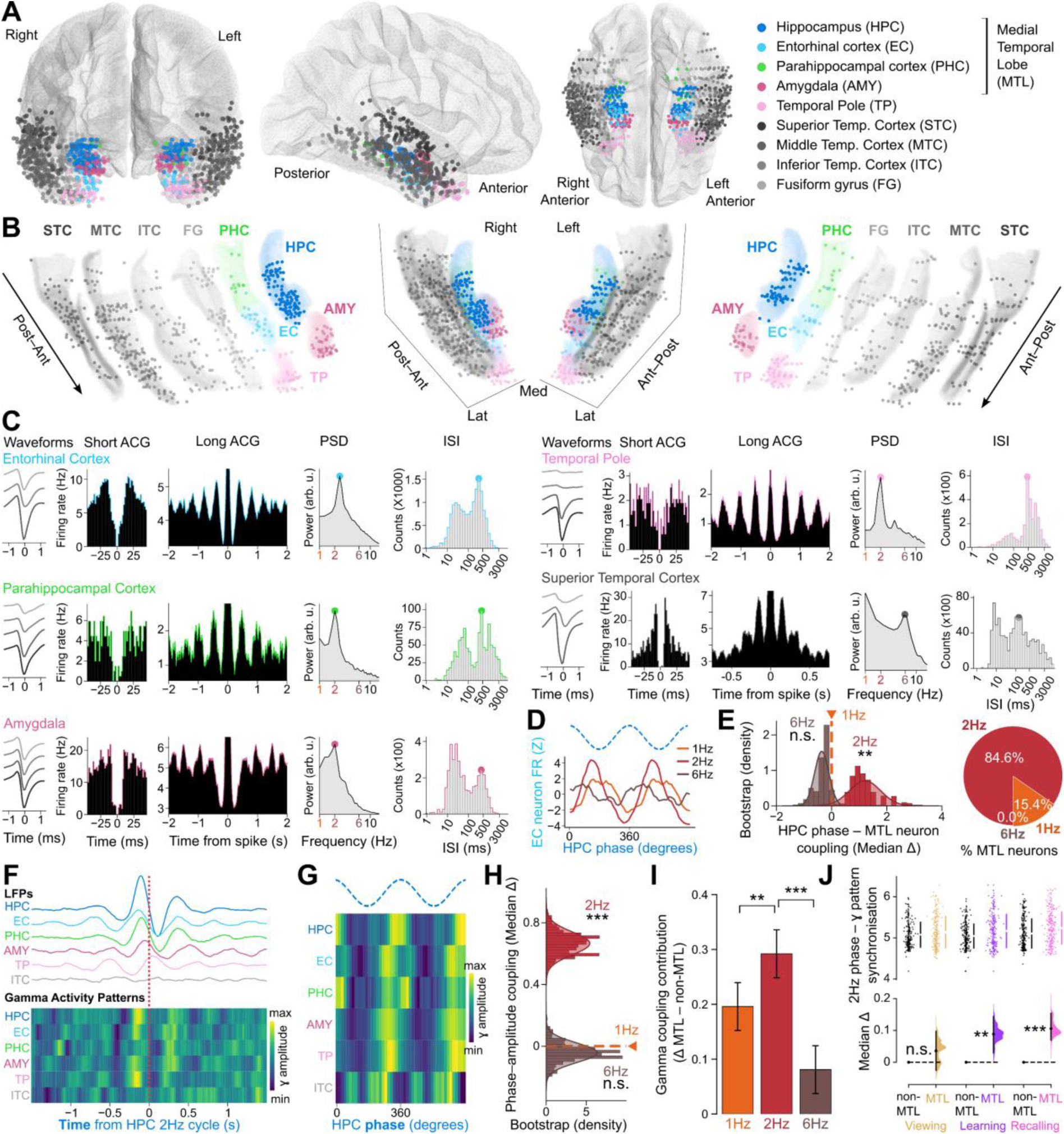
The hippocampal slow-oscillatory architecture synchronizes activity across the medial temporal lobe **(A,B)** MNI brain template **(A)** and 3D projection **(B**; with expanded view) showing electrode contact locations across MTL regions in all participants. **(C)** Representative neurons recorded from MTL tetrodes and exhibiting 2-Hz oscillatory rhythmicity, contrasted with a non-MTL neuron exhibiting faster (6-Hz) rhythmicity. **(D)** Firing-phase histograms of an example entorhinal cortex neuron relative to 1-, 2-, and 6-Hz hippocampal oscillations. **(E)** Median differences in spike–phase consistency between hippocampal 1-Hz and 2-or 6-Hz rhythms for extra-hippocampal MTL neurons and proportion of neurons preferentially coupled to each frequency. **(F)** Hippocampal slow oscillatory phase-triggered averages of LFPs from temporal lobe regions, showing rhythmic modulation of gamma-band activity. **(G)** Gamma amplitude averaged across hippocampal slow oscillatory phase bins. **(H)** Phase–amplitude coupling differences for MTL gamma envelopes relative to hippocampal 1-, 2-, and 6-Hz rhythms. **(I)** Difference in phase–amplitude coupling contributions (MTL minus non-MTL regression coefficients). **(J)** Estimation plot showing differences in slow oscillatory phase synchronization between MTL and non-MTL gamma activity patterns across task stages. Statistics were assessed using two-sided paired permutation tests or linear mixed models where appropriate. ***P < 0.001, **P < 0.01; n.s., not significant.

At the MTL network level, slow oscillatory power was most prominent in the hippocampus, whereas higher-frequency rhythmic activity dominated in extra-MTL regions (Figure S3A-C). The amplitude of 2-Hz oscillatory bursts did not correlate with ERP deflection in extra-hippocampal MTL regions (Figure S3D), a relationship that appeared specific to the hippocampus (Figure S1T). Importantly, hippocampal slow oscillatory phase synchronized gamma-band activity across MTL regions. Gamma activity patterns in extra-hippocampal MTL regions were more strongly coupled to hippocampal slow oscillatory phase (2-Hz) than to slower or faster rhythms (Figure 3F-H; 2-Hz versus 1-Hz, *P* < 0.001; 2-Hz versus 6-Hz, *P* < 0.001; 1-Hz versus 6-Hz, *P* = 0.320; two-sided paired permutation tests). Cross-frequency phase-amplitude coupling between hippocampal slow oscillatory activity and gamma patterns was stronger in MTL regions than in regions outside the MTL (Figure 3G,I; 2-Hz versus 1-Hz, *P* < 0.001; 2-Hz versus 6-Hz, *P* < 0.001; Wald tests on linear mixed-effects models).

This cross-regional synchronization was selectively enhanced during learning and recall compared to viewing (Figures 3J and S3E,F; viewing, *P* = 0.455; learning, *P* = 0.004; recalling, *P* < 0.001; two-sided paired permutation tests). Together, these results indicate that the hippocampal slow oscillatory architecture coordinates neural activity across the MTL, synchronizing distributed population dynamics during human memory processing.

### Cross-regional coactivity motifs structured by the transient coordination architecture during learning reactivate in hippocampal ripples

In animal models, population activity patterns formed during learning are later reactivated within hippocampal ripples to support memory consolidation during rest and sleep ^9,22,23,43–49^.

Hippocampal ripples also occur in humans ^50–55^, where they could similarly support post-learning consolidation. To test whether an offline reactivation mechanism operates in humans, we examined hippocampal ripple activity during pre-and post-learning rest sessions (Figures 4A and S4A-C).

**Figure 4.**
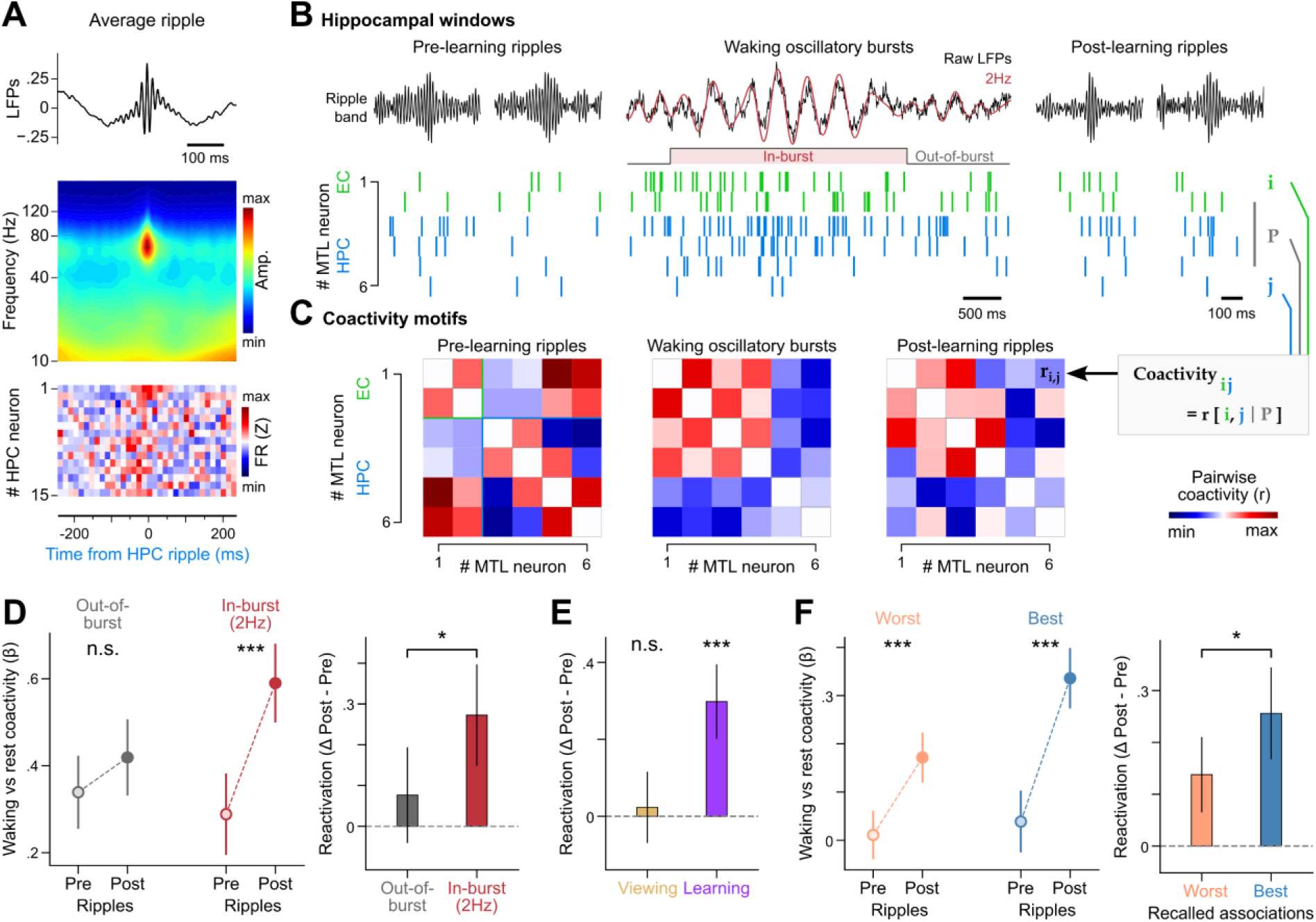
Cross-regional coactivity motifs organized by the hippocampal slow-oscillatory architecture are reactivated during offline ripples **(A)** Ripple-triggered average of hippocampal LFPs with corresponding spectrogram and firing-rate heatmap of hippocampal neurons. **(B,C)** Reactivation of waking (viewing and learning) MTL coactivity motifs during hippocampal ripples. MTL population spike trains were extracted from hippocampal ripples during pre-and post-learning rest, as well as from waking 2-Hz bursts (in-burst) and periods outside these bursts (out-of-burst) during viewing and learning sessions **(B)**. Coactivity motifs were derived from pairwise neuron-neuron (𝑖, 𝑗) correlations after regressing out global population activity 𝑃 **(C)**. **(D)** Similarity between waking and ripple coactivity motifs quantified by GLM β coefficients (left), and corresponding reactivation strength (right). **(E)** Selective reactivation of learning-related but not viewing-related coactivity motifs. **(F)** Reactivation of coactivity motifs detected during learning trials associated with best versus worst subsequent memory recall performance. Error bars indicate ±95% confidence intervals. Statistical significance assessed using two-sided Wald tests on GLM coefficients; ***P < 0.001, *P < 0.05; n.s., not significant.

The human hippocampus exhibited ripples that occurred reliably during rest and displayed spectral and temporal properties comparable to those recorded during sleep (Figure S4D-J). During rapid eye movement sleep, but not during slow-wave sleep, the human hippocampus also expressed slow oscillatory activity similar to that observed during waking task engagement (Figure S4K). To assess reactivation, we extracted MTL population spiking patterns expressed during hippocampal slow oscillatory bursts in viewing and learning sessions, and compared these waking “in-burst” patterns with population activity observed during hippocampal ripples in pre-and post-learning rest (Figure 4B). For comparison, we also extracted waking spiking patterns expressed outside slow oscillatory bursts and compared these “out-of-burst” patterns with ripple activity across rest sessions.

For each task session, we computed MTL coactivity motifs using pairwise neuron–neuron correlations while regressing out the activity of the remaining population (Figure 4C). Offline reactivation was quantified using generalized linear models predicting ripple coactivity from waking motifs, with interaction terms capturing changes in waking–ripple similarity between post-and pre-learning rest.

Following learning, coactivity motifs structured by the slow oscillatory architecture were selectively reactivated during hippocampal ripples. Specifically, ripples exhibited significantly stronger reactivation for coactivity motifs expressed during slow oscillatory bursts than for those expressed outside these bursts (Figure 4D; in-burst, *P* < 0.001; out-of-burst, *P* = 0.202; in-burst versus out-of-burst, *P* = 0.025; Wald tests), or for motifs associated with slower or faster rhythmic activity (Figure S4L). Moreover, population coactivity evoked by photographic stimuli during learning reactivated more strongly during post-learning ripples than that evoked during viewing (Figures 4E and S4M-O; learning, *P* < 0.001; viewing, *P* = 0.622; learning versus viewing, *P* < 0.001; Wald tests). Critically, coactivity motifs associated with the best subsequent memory performance showed the strongest post-learning ripple reactivation (Figure 4F; best and worst recalled, both *Ps* < 0.001; best versus worst recalled, *P* = 0.043; Wald tests).

Together, these findings demonstrate that population dynamics structured by the slow oscillatory architecture during learning are selectively reactivated during hippocampal ripples, linking online memory processing to offline consolidation in humans.

## Discussion

Our findings establish a unifying organizational principle for human memory in which slow oscillatory events provide the temporal backbone that links online encoding, offline consolidation, and subsequent recall across the medial temporal lobe. This architecture organizes population spiking, synchronizes gamma-band activity across hippocampal and extra-hippocampal regions, and structures patterns of neuronal coactivity that are selectively reactivated during offline hippocampal ripple events. Rather than reflecting a continuous background rhythm, it operates through transient, task-evoked events that define temporal windows for coordinated network interactions during memory processing.

Human hippocampal oscillatory activity has long appeared variable^37,38^, leaving unresolved whether oscillatory dynamics in the MTL provide an organizing framework for memory comparable to those described in animal models. Intracranial recordings have reported slow-frequency oscillations in the human hippocampus, including activity in the ∼2–4 Hz range associated with memory encoding and gamma coupling, whereas effects in the conventional rodent theta range are often weaker or variable^34–36,42^. Other studies have nonetheless linked theta-range power, entorhinal stimulation, thalamic stimulation, and cholinergic modulation to memory performance, highlighting the behavioral relevance of slow-frequency dynamics in human memory circuits^56–59^. Our results provide a mechanistic framework for interpreting these diverse observations by identifying a slow oscillatory architecture that organizes MTL activity during memory processing in humans.

In animal models, hippocampal oscillations provide a temporal framework for coordinating memory-related neural activity. In rodents, theta-band oscillations organize population spiking during active behavior, support interactions with nested gamma-band activity, and structure patterns of neuronal coactivity during learning^14,17–20^. These patterns are subsequently reactivated during hippocampal ripples in rest and sleep, supporting offline consolidation^9,22,23,43–49^. Together, these findings have shaped influential models of memory and cognition in which oscillatory dynamics organize mnemonic processing, including frameworks that link online encoding and recall with offline consolidation^7,10,26–29^.

However, accumulating evidence indicates that hippocampal oscillatory dynamics vary substantially across species and behavioral contexts. In larger-brained mammals, including rabbits, cats, bats, and primates, hippocampal rhythmic activity is often slower, more intermittent, and more closely linked to task demands than to locomotion^30–33^. In humans, reports of MTL oscillatory activity have emphasized its variability and intermittent nature^37,38^, raising the question of whether such activity can provide an organizing architecture capable of coordinating neuronal spiking, network synchronization, and offline reactivation across memory states.

Our findings address this gap by demonstrating that despite their transient nature, ∼2-Hz bursts anchored in the hippocampus structure memory-related network dynamics across the human MTL. These bursts pace population spiking, synchronize gamma-band activity across hippocampal and extra-hippocampal regions, and structure coactivity motifs that are selectively reactivated during offline hippocampal ripple events, with reactivation strength predicting subsequent recall accuracy. Notably, the coordinating effects we observe are selective to slow oscillatory activity centered near 2-Hz and are not present at slower (1-Hz) or faster (6-Hz) frequencies, indicating that this architecture operates at a constrained organizing timescale rather than reflecting broadband slow fluctuations. Through this frequency-specific organization of network interactions, oscillatory coordination during learning is directly linked to offline consolidation and later memory performance.

This slow oscillatory architecture can be interpreted as a species-adapted organizing mechanism that preserves the logic of oscillatory coordination while operating at a slower timescale in the human brain. As brains enlarge across species, maintaining coherent timing among distributed circuits becomes increasingly challenging. Compensatory adaptations such as increased axon caliber and myelination help offset conduction delays introduced by longer conduction paths, preserving relative temporal relationships across regions ^60^. Within this context, a slower coordinating rhythm may facilitate communication across distributed networks without compromising faster nested dynamics for local computation. Consistent with this scaling framework, the peak frequency of the hippocampal theta rhythm, long implicated in waking memory processing in animal models, decreases with brain size across mammals, from ∼5–12 Hz in mice and rats to ∼3–7 Hz in bats, rabbits, cats, and primates ^10,16,30–33,61,62^. The identification of a slow organizing rhythm around ∼2 Hz that paces online memory dynamics in humans extends this continuum without implying strict frequency homology. Moreover, hippocampal expression of this slow oscillatory architecture during rapid eye movement sleep^63,64^, further supports the idea that a conserved organizing logic operates across brain states as well as across species.

Importantly, our findings do not suggest that memory coordination in humans depends on a direct analogue of rodent theta. Instead, they support a conserved organizing principle in which slow oscillatory dynamics structure population activity, coordinate cross-regional communication, and link online encoding with offline consolidation. Together, these results show that human memory is organized by a scalable oscillatory architecture that coordinates network dynamics across learning, consolidation, and recall through mnemonic engagement-locked events. Without this transient, slow oscillatory architecture, a core principle of memory-related network organization in humans linking neuronal spiking, cross-regional communication, and ripple-mediated consolidation would remain mechanistically unexplained.

## Methods

### Subjects

This study included a total of 35 adult participants [mean age (interquartile range): 35 (24–42) years; 16 males, 19 females; 26 right-handed] undergoing intracranial monitoring for pharmacologically intractable epilepsy. Participants were recruited from the Epilepsy and Sleep Unit of the Neurology Department at Toulouse University Hospital (Toulouse, France; n = 24) and from the Epilepsy and EEG Units at Pitié-Salpêtrière Hospital (Paris, France; n = 11). All patients underwent stereo-electroencephalography with multi-contact depth electrodes over 7–10 days to localize seizure foci. Electrode placement followed clinical requirements and pre-surgical trajectories determined from individual structural MRI scans. All participants provided informed consents following the procedures approved by the relevant institutional review boards.

### Intracranial electrodes

Participants were implanted with standard and hybrid depth electrodes. In Toulouse, hybrid electrodes (DIXI Medical; maximum of six electrodes per subject; platinium/iridium, 2 mm in length, 0.8 mm in diameter) comprised 5–18 macrocontacts and were equipped with two or three tetrodes (tungsten wires, 20 μm in diameter), as described in ^41^.

Tetrodes typically protruded between the two deepest macrocontacts on each electrode (Figure 1E); in few hybrid electrodes, they extended between the eighth and the ninth macrocontacts to sample more superficial regions. The reference for both tetrodes and macrocontacts was a macrocontact located in white matter. Bilateral implantations allowed recording from contralateral, unaffected sites (sentinel electrodes). In Paris, hybrid electrodes contained 9 macrocontacts (3 mm spacing between contact 1 and 2, and 4.5 mm for the others) and eight microwires (Behnke-Fried electrodes, Ad-Tech Medical; maximum of two electrodes per subject), with one additional microwire used as a reference. Standard electrodes had 5 mm spacing between contacts. The anatomical location of each macrocontact was determined by co-registering postoperative CT scans with preoperative 3D T1-weighted MRI data, as presented in^65^. Preoperative T1 3T MRI images were automatically segmented in the native space (using FreeSurfer, https://surfer.nmr.mgh.harvard.edu/), and the anatomical location of each electrode macrocontact was then detected on the CT scans and positioned in the Desikan-Killiani atlas ^66^. The location of all gray matter contacts and Behnke-Fried microwire bundles were further verified by an expert clinician. Each tetrode was assigned the anatomical label of the nearest macrocontact. To localize macrocontacts within the hippocampal volume, each macrocontact was assigned a standardized anatomical position based on individual hippocampal geometry. Hippocampal volumes were segmented from native T1-weighted MRI scans, and segmentation quality was manually verified (used in Figures S1A,J, S2A, S3A). All contact coordinates were then normalized in the Montreal Neurological Institute (MNI) space for group-level analyses and visualizations (Figures 1F,G, 3A,B). Hippocampal macrocontacts with y-axis coordinates <-22 in the MNI space were considered posterior (Figure S2E). Of 170 contacts, 105 were located in the right hemisphere.

### Neurophysiological recordings

Electrophysiological signals were recorded continuously for approximately one hour while participants performed the behavioral task, using an ATLAS 16SX 256-channel Neuralynx. For overnight sleep, additional surface electrodes were also used according to the 10–20 system alongside electrooculography and electromyography electrodes for polysomnography ^67^. Data were filtered between 0.1 and 8,000 Hz and sampled at 32,768 Hz, then downsampled to 20,000 Hz and 1,250 Hz for subsequent analyses of population spiking and neural oscillations, respectively. Macrocontact and microelectrode signals were visually inspected to exclude artifactual channels. In total, 2,667 macrocontacts were retained (99 ± 24 per subject). A common-average referencing was applied to macrocontact data (using the median to minimize the potential influence of large-amplitude events including interictal epileptiform discharges) but not to microelectrode recordings. Where mentioned in the manuscript, bipolar referencing was occasionally used for control analyses (Figure S1H,I). Line noise at 50 Hz was minimal under the recording conditions; consequently, notch filtering was never applied.

Synchronization between stimulus events (e.g., photographic presentations) and neural recordings was achieved via TTL pulses, with each event type assigned a unique TTL identification code. Throughout the manuscript, local field potentials (LFPs) refer to intracranial EEG signals recorded from macrocontacts, except in Figure 2A and S4B,C where tetrode LFPs were used. LFPs from microwires (Behnke-Fried electrodes) are not used in this study because local referencing was employed for initial acquisition of the signal. Consequently, Behnke-Fried electrodes are not displayed in the figures of this manuscript.

### Behavioral datasets

Twenty-seven participants (n = 16 in Toulouse; n = 11 in Paris) performed a computer-based associative memory task implemented in MATLAB using Psychophysics Toolbox 3 (PTB-3). The task involved learning associations among individuals forming a community, in which each person was represented by a photograph and a name. Each recording day consisted of five consecutive sessions: pre-learning rest, viewing, learning, post-learning rest, and recall (Figure 1A; total duration: ∼1 hour). During the two rest sessions, participants sat still with eyes closed (10 minutes each session). In the viewing session (8 minutes), they were familiarized with 144 photographs of community members displayed for 1 second each and in random order, separated by 1.5–3 second inter-stimulus intervals sampled from a gamma distribution, during which a central fixation dot was shown. Participants were instructed to maintain fixation. To ensure compliance, we asked them to press the space bar of the computer keyboard whenever they detected that the same photograph was presented consecutively, a repetition that we introduced at random intervals. We then briefly familiarized participants to the structure of the learning session (1–2 minutes). During the learning session (10 minutes), participants viewed pairs of individuals presented consecutively, one person after the next, to associate them (i.e., “paired associates”). Between each pair, participants were instructed to maintain fixation on the displayed central dot. Learning progress was assessed through intermittent multiple-choice questions: a cue photograph appeared for 1 second, followed by a screen displaying three possible associates. Participants had up to 10 seconds to identify the correct associate and respond using the left, down, or right arrow keys. In this community, one individual was intermittently shown holding a key to a treasure chest and participants were asked to remember it. After the post-learning rest session, participants performed the recall session (15 minutes) to test retention of the learned associations. During recall, participants answered two types of memory questions. The first type consisted of multiple-choice questions identical to those used during learning, in which participants identified the correct associate among three photographs. The second type presented a single photograph for 1 second, after which participants judged whether the depicted person could help obtain the key based on the associations they had learned. Responses were permitted after a 2-second delay, creating a fixed 3-second interval between cue onset and the earliest possible motor response. This delay ensured that the visual display was directly comparable to the viewing and learning photographs for analysis of event-related potentials and evoked oscillations (Figures 1J,K and S1S-U). The recording day finished with a post-recall viewing session, which we used to confirm that the changes in oscillation power during the memory recall session were not merely reflecting passage of time (data not shown).

We further validated the network physiology identified in the awake human hippocampus during the relational memory task by examining data from eight additional awake participants who were watching a sitcom series (*Friends*), and were instructed to maintain attention throughout each episode (∼25 minutes per episode). A member of the team continuously monitored participants’ vigilance to ensure sustained attention.

To further validate our ripple detection in the human hippocampus during rest sessions of the relational memory task, we also characterized oscillatory patterns expressed during overnight sleep in seven of our participants recruited to do the memory task. These participants were recorded continuously for 24 hours including overnight sleep. Sleep staging was performed according to American Academy of Sleep Medicine guidelines ^67^. N2 and N3 stages were merged and referred to as slow-wave sleep throughout the manuscript.

### Spike detection and unit isolation

Spike sorting and unit isolation were performed with an automated clustering pipeline using Kilosort (https://github.com/cortex-lab/KiloSort) via the SpikeForest and SpikeInterface frameworks (https://github.com/flatironinstitute/spikeforest) ^68,69^, as previously used in rodent studies ^39,70^. For this, the algorithm restricted templates to channels within a given microelectrode while masking all other recording channels. All sessions recorded contiguously on a given day were concatenated and cluster cut together to monitor cells throughout the experiment. The resulting clusters were verified by the operator using cross-channel spike waveforms, auto-correlation histograms, and cross-correlation histograms. Each unit used for analyses showed throughout the entire recording day stable spike waveforms, clear refractory period in their auto-correlation histogram, and absence of refractory period in its cross-correlation histograms with the other units. All analyses were performed on these single units. In total, 257 single units were identified in the MTL, of which 211 in the hippocampus (Figure S2L). All hippocampal single-units analyzed in this manuscript were detected using tetrodes (from n = 18 human participants out of the 24 recruited in Toulouse). This decision was made to ensure that hippocampal neurons were located close to the macrocontacts used for LFPs extraction (e.g., to detect the phase of local oscillations; see model of the hybrid electrode in Figure 1E). In the extra-hippocampal MTL, 39 single units were detected with tetrodes, and 7 using microwires (Behnke-Fried electrodes).

### Behavioral analysis

Behavioral performance during learning and memory recall was quantified as the percentage of correct responses to multiple-choice questions. This was assessed intermittently across trials during the learning session (Figure 1B) and as the mean accuracy across all questions during the recall session (Figure 1C, Wilcoxon signed-rank test). To identify the best and worst recalled associations (Figures 1K, 4F), accuracy was calculated for each individual association within the learnt community. For each association, the mean accuracy across memory questions was computed. The two associations with the highest mean accuracy were labeled as “best”, and the two with the lowest mean accuracy as “worst”. Recording days when subjects exhibited ceiling performance across all associations were excluded from this analysis (n = 2 out of 34 recording days excluded).

### Decomposition of LFPs into oscillatory components

To decompose the local field potentials (LFPs) into their elementary oscillatory components, or intrinsic mode functions (IMFs), we applied tailored masked Empirical Mode Decomposition (tmEMD) ^40^. EMD is an unsupervised, iterative sifting algorithm that extracts signal components based on local time-frequency properties. Masked EMD (mEMD) enhances this process by introducing a mask signal at each sifting step to reduce mode mixing and thus improve the separation of IMFs. The tailored variant of mEMD was developed to minimize mode mixing, increasing consistency across recordings from different subjects and electrode contacts. Using the macrocontacts located in the anatomical region of interest (e.g., hippocampus in Figure S1D-F), we applied the tmEMD procedure to LFP recordings using two steps. The first step allowed assessing the epochs to be analyzed and optimizing the masks. For each signal, we used 2 × 45-s LFPs epochs free of interictal epileptiform discharges (IEDs) and with high signal-to-noise ratio (SNR). To quantify SNR for a given epoch, we computed the Welch PSD and calculated the power ratio between 1–20 Hz and >200 Hz. Mask frequencies were optimized by pooling the identified clean epochs across macrocontacts and applying tmEMD to identify mask parameters that maximize consistency and minimize mode mixing. The second step performed the full-session decomposition using optimized masks: once optimal masks were identified, we applied mEMD to the full recording session. Instantaneous amplitude, frequency, and phase were then obtained using the Hilbert transform. IMF consistency was assessed by computing the PSDs of each IMF in IED-free epochs (Figure S1D,F). To characterize the frequency band of each IMF, we computed its PSD using Welch’s method and averaged the resulting spectra across all macrocontacts (Figure S1E thick lines). For each IMF, the peak frequency was defined as the frequency of maximum power, and the frequency band was determined as the contiguous band containing 80% of the total spectral power (i.e., 80% power band).

### Power spectral densities

Power spectral density (PSD) estimates were computed using Welch’s method with a Hann window (from the scipy.signal.welch function in the scipy.signal module). PSDs were calculated across contiguous 1-s windows with a frequency-dependent sampling density optimized for slow and fast oscillations (20 points/Hz for 0.5–20 Hz; 4 points/Hz for 20–200 Hz). All LFPs signals were z-scored prior to spectral estimation (Figures 1H and S4B,K). To account for interindividual differences in the broadband (aperiodic) component of the power spectrum, we modeled each PSD using the “Fitting Oscillations and One-Over-F” (FOOOF) algorithm ^71^.^71^. Each PSD was fit within the 0.5–20 Hz range to isolate the aperiodic (1/f) component. The full modeled spectrum and the aperiodic fit were then subtracted to obtain a log-corrected, aperiodic-adjusted PSD, preserving only periodic features (Figures S1I,K–N and S3A–C). Narrowband oscillations were then quantified by averaging power within the 1-Hz (0.5–1.25 Hz), 2-Hz (1.25–3.5 Hz), and 6-Hz (3.75–8.5 Hz) frequency bands, corresponding to the power distribution of the initially detected IMFs (Figure S1E). In addition, to characterize rhythmicity in single-neuron spiking activity, PSDs were computed on spike-train autocorrelograms. For each unit, the autocorrelogram was calculated over a 2-s window and smoothed with a Gaussian kernel (width = 10 ms, σ = 5 ms) to reduce high-frequency noise while preserving rhythmic structure. The PSD of the resulting autocorrelation signal was then computed (0.9–15 Hz range) to identify oscillatory modulation of firing patterns. Peaks in these autocorrelogram-derived PSDs indicate dominant rhythmic frequencies in the spike timing of individual neurons (Figures 2F, 3C and S2M).

### Detection of interictal epileptiform discharges (IEDs)

We detected IEDs recorded in our datasets using an envelope-based, data-driven algorithm adapted from Janca *et al.* ^72^.^72^. The continuous LFPs traces were downsampled to 200 Hz and band-pass filtered (Chebyshev type-II high-pass 10 Hz and low-pass 60 Hz, stopband attenuation 30 dB). The analytic envelope (Hilbert) was computed and scanned with overlapping 5-s windows stepped by 1 second. Within each window, the envelope distribution was modeled by a log-normal function, and events were detected when the signal amplitude exceeded three times the typical (mode + median) level estimated from that distribution. This adaptive thresholding identifies transient amplitude outliers that deviate from the typical background activity. Window thresholds were cubic-spline interpolated to sample resolution and boxcar-smoothed, and contiguous supra-threshold samples were grouped into events. Nearby events were merged by extending boundaries by ±120 ms and the envelope peak within each merged window was taken as the IED time. All detections were produced on the downsampled signal and indices were mapped back to the native sampling rate for downstream analyses. Examples of detected events are shown in Figure S1J. Macrocontacts showing an average detection rate below one IED per minute during active sessions were considered free of epileptiform activity.

### Comparison of oscillatory power between task sessions

Session-related changes in oscillatory power were quantified across task sessions for the 1-, 2-, and 6-Hz frequency bands. Power values were obtained from log-corrected PSDs computed for each hippocampal macrocontact as described above (see section “*Power spectral densities*”), excluding those with interictal activity (Figure S1L–N). Linear mixed-effects models were used to compare power across sessions, with task session treated as a fixed effect and individual subject as a random factor, allowing assessment of frequency-specific modulation of slow oscillations throughout the task, controlling for putative subject-driven effects (Figure S1N).

### Detection of oscillatory bursts

Oscillatory bursts were detected from IMF-derived signals using a spectrogram-based thresholding approach. For each hippocampal recording, the signal reconstructed up to the IMF of interest (1-, 2-or 6-Hz IMF) was converted to a wavelet spectrogram using 30 logarithmically spaced (0.2–20 Hz for 1-Hz bursts, 0.5–25 Hz for 2-Hz bursts, and 1–45 Hz for 6-Hz bursts). The spectrogram was z-scored, and contiguous time-frequency clusters exceeding a z-score of 2 were identified as candidate bursts. Clusters were retained only if at least half of their energy fell within the target frequency band (0.5–1.5, 1–4, 5–12 Hz for 1-, 2-and 6-Hz bursts, respectively) and if their duration corresponded to at least two oscillatory cycles. Adjacent clusters separated by <400 ms (1-Hz), <200 ms (2-Hz) or <50 ms (6-Hz) were merged. Example 2-Hz bursts are shown in Figure S1O. Burst onset and offset were then defined from the merged clusters, and each burst’s mean frequency, duration, and cycle count were extracted (Figure S1P,Q). Out-of-burst epochs used in the reactivation analyses (Figures 4D) corresponded to all segments between detected bursts.

### Comparison of event-related potentials between sessions

Event-related potentials (ERPs) were measured by averaging LFPs time-locked to photograph onsets. For each recording, signals were z-scored, and trials contaminated by IEDs or falling outside the recordings were excluded. Epochs extending from 1 second before to 4 seconds after photograph onset were extracted, baseline-corrected using the mean pre-stimulus activity, and averaged across trials for each contact. To test for session-related differences, the two average ERPs were compared using a nonparametric cluster-based permutation test (1,024 iterations, mne-python package ^73^), which identifies contiguous time intervals showing consistent ERP deflections differences while controlling for multiple comparisons. Significant clusters (p < 0.01) indicated time windows with reliable modulation of evoked responses across sessions (Figure S2R).

### Wavelet spectrograms and quantification of evoked oscillations

Amplitude spectrograms were computed from the raw signal using complex Morlet wavelet convolution for each macrocontact across sessions. This approach provides high temporal precision for fast frequencies and high frequency resolution for slow oscillations, making it well suited to characterize non-stationary neural dynamics. Wavelets were constructed with five cycles per frequency and normalized to unit energy, producing 80 logarithmically spaced frequencies between 1 and 150 Hz. Stimulus-locked spectrograms were then extracted from –1 to +3 seconds (viewing and recalling sessions) or +4 seconds (learning session) relative to stimulus onset, excluding trials containing IEDs or motor responses. To ensure comparability across sessions, trials were subsampled to equalize stimulus counts. For each contact and frequency, post-stimulus amplitudes were normalized to baseline (–1 to 0 second) by z-scoring and averaged across trials using the amplitude from the baseline only (Figures 1J, S1T,U and S3D). This baseline normalization was not applied in Figure S1S, where the mean amplitudes in the post-stimulus window were compared to mean amplitudes obtained from the baseline. Mean evoked amplitudes were then quantified within the 1-, 2-, and 6-Hz frequency bands during the post-stimulus window (+1 to +3 seconds for viewing and recalling, +2 to +4 seconds for learning).

This ensures that the observed changes in evoked amplitude are not confounded by differences in the magnitude of ERPs deflection between conditions.

### Correlation between ERP deflection and evoked oscillatory amplitude

To relate ERPs deflection with evoked oscillations in the recalling session, we compared the mean ERP deflection (0.3–0.9 second after stimulus onset in the hippocampus) with the corresponding post-stimulus spectral amplitude averaged within the 1-, 2-, or 6-Hz bands (1–4 second window).

Correlations were computed across contacts using Pearson’s coefficient, and significance was assessed by two-tailed testing. Scatterplots in Figures S1T and S3D display the linear fit with bootstrapped 95% confidence intervals, showing the relationship between the strength of evoked slow oscillations and the magnitude of the ERP deflection. The mean ERP deflection was taken between 0.3–0.9, 0.15–0.6 and 0.15–0.5 second after stimulus onset for the entorhinal cortex, parahippocampal cortex and amygdala macrocontacts, respectively (Figure S3D). These epochs were selected to match the biggest ERP deflection visible in the grand averages computed for each region. However, various epochs were tested and none of them showed significant correlation with evoked amplitude (*data not shown*).

### Detection of gamma activity

To detect gamma activity, the signal was divided into five contiguous frequency bands between 60 and 160 Hz (20-Hz steps). Each band was bandpass-filtered and transformed into its analytic representation using the Hilbert transform to obtain the instantaneous amplitude. The envelope from each band was normalized by its mean amplitude to equalize their contributions, and the normalized envelopes were averaged across all bands to produce a single broadband gamma envelope representing moment-to-moment fluctuations in gamma activity (Figure 2A).

### Detection of individual oscillatory cycles

To segment individual oscillatory cycles from each intrinsic mode function (IMF), we identified four key waveform features: troughs (local minima), peaks (local maxima), ascending and descending zero-crossings. Cycles were defined as sequences of six temporally ordered points: two consecutive troughs (or alternatively, two peaks, or two zero-crossings), and four intermediate landmarks corresponding to characteristic inflection points within the waveform (e.g., zero-crossings and extrema). Each cycle was retained only if all six points followed a strictly increasing temporal order, ensuring well-formed and physiologically plausible waveforms. Additional checks were applied to exclude cycles with overlapping or missing components. This procedure enabled the consistent identification of complete cycles across IMFs (Figures 2B, 3F, 4B, and S2A,C).

### Oscillatory cycle-triggered averages

To relate slow oscillatory phase (1-, 2-, and 6-Hz) to gamma activity and population firing rate (Figures 2B and S2C), we computed cycle-triggered averages aligned to each extracted IMF cycle. Cycles that coincided with IEDs were excluded. For each cycle, we extracted ±2.5 seconds around the descending zero-crossing and binned signals at 10, 5, or 3 ms resolution for 1-, 2-, and 6-Hz cycles, respectively. LFPs were z-scored, and spike counts were converted to population rate (Hz) per bin. Cycle-averaged LFP, gamma, spike-rate traces, and time–frequency spectrograms (>10 Hz) were obtained by averaging across cycles (and across units for the population rate). To visually compare the three frequencies, gamma activity was normalized, and the same y-scales were applied to all graphs (Figures 2B and S2C).

### Phase–amplitude coupling and spike–field locking

To quantify the actual phase–amplitude coupling (PAC) between gamma activity and slower oscillations, we computed the modulation index (MI) as in ^74^. Gamma signals were taken either from the same macrocontact signal as the phase (local PAC; Figures 2E and S2E,F) or from a distal macrocontact (distal PAC, across structures; Figure 3H,I). The computation of the MI involved measuring the deviation of the amplitude distribution across phase bins from uniformity using the Kullback–Leibler divergence. First, we measured the instantaneous phase of the IMFs using the Hilbert transform. The phase of each IMF (e.g., 2-Hz) was then binned into *N* equally spaced intervals over [0, 2π]. For each bin *i*, we computed the mean amplitude of the gamma activity *A[n]* as:

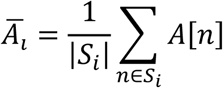

where 𝑆_𝑖_ is the set of time points for which *ϕ[n]* falls into bin *i*. We then normalized the amplitude distribution:

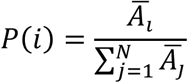

The Kullback–Leibler divergence between this distribution and the uniform distribution 𝑈(𝑖) = 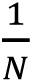 was computed as:

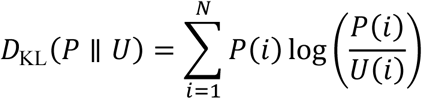

We normalized this value to define the modulation index:

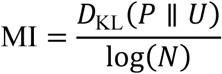

This yielded a value between 0 (no coupling) and 1 (maximum coupling), indicating how strongly the amplitude of the gamma activity depends on the phase of the slow oscillation. Then, we generated 300 surrogate versions of the intrinsic mode functions (IMFs) while preserving their spectral content by applying phase randomization in the Fourier domain. This approach destroys temporal structure but retains the power spectrum, making it suitable for surrogate-based statistical testing of the PAC.

The original signal *x[n]* was transformed using the discrete Fourier transform (DFT):

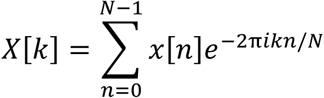

where *A[k]* is the amplitude spectrum and *ϕ[k]* the phase spectrum. This was expressed in polar form as:

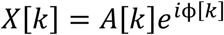

where 𝐴[𝑘] = |𝑋[𝑘]| is the amplitude spectrum, and 𝜙[𝑘] = arg(𝑋[𝑘]) is the phase spectrum. We then generated a random permutation *π(k)* over frequency indices and substituted:

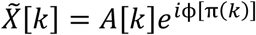

The inverse Fourier transform yielded the phase-randomized surrogate signal:

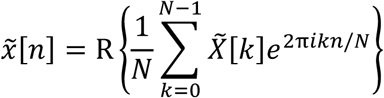

We then computed phase–amplitude coupling (PAC) between the unchanged gamma activity and each of the 300 surrogate phase signals. This generated a null distribution of PAC values, which we used to assess statistical significance. The actual PAC value, computed using the original (non-randomized) phase signal, was transformed into a z-score relative to this surrogate distribution. This z-scored PAC reflects the likelihood—rather than the magnitude—of coupling and was used for statistical comparisons across IMFs and electrode contacts (Figures 2E, 3H and S2E).

To assess the relationship between spike timing and the phase of slower oscillations, we computed the pairwise phase consistency (PPC) as in ^75^. Spike trains were either taken from neurons located in the same region as the phase (local PPC, nearest macrocontact; Figure 2C,D) or from a distal region (distal PPC, across structures; Figure 3D,E). For each neuron, we extracted the instantaneous phase of the IMF (e.g., 2-Hz) at the time of spike occurrence. Given a set of *N* spike times, each associated with a phase value 𝜙_𝑛_ ∈ [0, 2𝜋] the PPC is computed as:

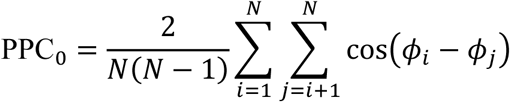

This expression corresponds to the average cosine of the phase differences between all unique spike pairs. It can also be reformulated for computational efficiency using trigonometric identities:

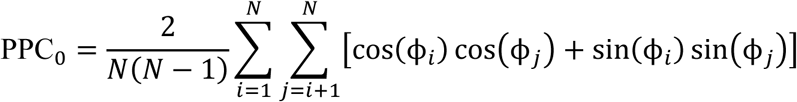

The PPC value ranges from 0 (uniform/random phase distribution) to 1 (perfect phase locking). To determine whether the observed spike–phase coupling was statistically significant, we used the same phase randomization followed by z-scoring procedure described above for PAC. Specifically, we generated 300 surrogate phase signals by shuffling the phase time series and recomputed the PPC between each surrogate phase and the spike times. This yielded a null distribution, against which the PPC computed from the original phase signal was z-scored. This z-scored PPC reflects the likelihood of spike–phase locking beyond chance and was used for statistical testing across IMFs and neuronal units (Figures 2C,D and 3D,E). A z-score threshold of 5 was used to identify significant coupling. Preference was then measured by comparing, for each given neuron, which of the three oscillations (1-, 2-, or 6-Hz) had the highest z-scored PPC value, only considering neurons with at least one significant locking in these frequency bands (Figure 3E, left). Both PAC and PPC were measured on epochs clear from interictal discharges.

### Quantification of phase reversal

Laminar phase relationships along depth electrode contacts were analyzed to identify polarity reversals between adjacent hippocampal recording sites.

Average LFPs were first computed from three neighboring macrocontacts, aligned to the positive peaks of 2-Hz cycles detected on the deepest contact (e.g., “Hpc 1a” in Figure S2A). The 2-Hz instantaneous phase was then extracted from IMFs on each macrocontact using the Hilbert transform, and circular statistics were applied to determine the mean phase difference between all contact pairs (Figure S2B).

### Cross-correlation between gamma activity and population firing rate

To assess the temporal coupling between local neuronal firing and gamma activity, we computed time-lagged cross-correlations between population rate and gamma activity. For each recording session, we retained hybrid electrodes located in the hippocampus and with at least five recorded neurons.

Population rate was obtained by summing spike trains from all single-units recorded within the same region and temporally smoothing the resulting spike-rate vector with a Gaussian kernel (width = 50 ms, σ = 25 ms). For each region, Pearson correlation coefficients were computed between the population rate and each local gamma trace across contiguous, IEDs-free epochs. To evaluate specificity, the same population rate signal was correlated with gamma envelopes from distal macrocontacts located on other electrode shafts. Mean correlation coefficients across distal contacts were used as controls (Figure S2C).

### Anatomical gradients in local phase–amplitude coupling

Anatomical gradients in local phase–amplitude coupling (PAC) were assessed to determine whether the strength of oscillatory coupling varied along hippocampal axes. For each hippocampal contact, coupling values were related to the anteroposterior and mediolateral coordinates of this recording site using a generalized linear model fitted separately for the 1-, 2-, and 6-Hz frequency bands. Coupling values were transformed to match normality of the distributions (Yeo-Johnson):

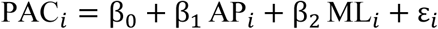

where PAC_𝑖_ is the transformed local phase–amplitude coupling value for contact 𝑖; AP_𝑖_ and ML_𝑖_ are the anteroposterior and mediolateral coordinates of this recording site; 𝛽_0_ is the intercept; 𝛽_1_and 𝛽_2_ represent the effects of position along the hippocampal axes; and 𝜀_𝑖_ is the residual error term. Model coefficients were extracted and summarized as a heatmap (Figure S2F).

### Amplitude modulation using Holo-Hilbert Spectral Analysis (HHSA)

The HHSA is designed to analyze non-linear and non-stationary signals and capture both carrier frequencies and their amplitude modulations ^76^. This spectral method uses a two-layer EMD followed by Hilbert transforms to provide a two-dimensional representation of energy across carrier frequencies and modulation frequencies. This allows dealing with non-linearities and harmonics and reduces the risk of detecting spurious PAC ^77^ to identify genuine cross-frequency interactions and nested oscillations in LFPs signals. To do this, we first selected epochs clear from interictal epileptic discharges and used IMFs extracted using tmEMD on each macrocontact, retaining the first *K=9* IMFs capturing slow to fast oscillations. Each IMF was transformed into its analytic signal via the Hilbert transform:

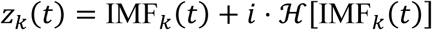

yielding instantaneous amplitude 𝐴_𝑘_(𝑡) = |𝑧_𝑘_(𝑡)| and carrier frequency:

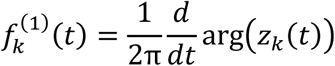

To capture amplitude fluctuations over time, we applied a second-layer EMD to the amplitude envelopes, decomposing each into slower amplitude modulation components:

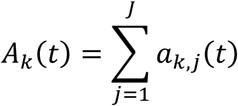

Each modulation component was again Hilbert-transformed to extract modulation frequency. The holospectrum represents signal energy jointly as a function of carrier frequency 𝑓^(1)^ and amplitude modulation frequency f ^(2)^. It was computed across all IMF pairs and projected onto a 2D frequency space:

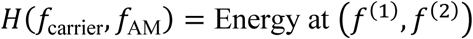

Log-spaced frequency bins were used from 0.1 to 195 Hz for both axes, allowing fine resolution of slow-frequency modulations. The average energy measured on each electrode contact signals were then z-scored before averaging across contacts. We finally averaged the energy in several frequency bands to compare gamma (60–160 Hz) modulation across slow oscillations (Figure S2G–K) That is, in humans, 1-Hz (0.5–1.25 Hz), 2-Hz (1.25–3.5Hz), and 6-Hz (3.75–8.5 Hz); in mice, 3-Hz (1.5–4.5 Hz), 7-Hz (5–10 Hz), and 15-Hz (11–20 Hz). To quantify the gain in amplitude modulation in contacts clear from interictal discharges, we computed the paired difference between 2-Hz and 6-Hz energy per macrocontact (Figure S2I).

### Comparison of oscillatory power between brain regions

Regional differences in oscillatory power were assessed by comparing log-corrected spectra between hippocampal (HPC), extra-hippocampal medial temporal (MTL; entorhinal, parahippocampal, amygdala), and non-MTL temporal cortical contacts (inferior, middle and superior temporal cortices and fusiform gyrus). For each macrocontact, log-corrected spectra were computed as described in the Methods section *“Power spectral densities”* (Figure S3A). To better capture the dominance of alpha oscillations over 2-Hz in non-MTL regions, we then used the peak power measured as the maximum of the corrected power spectrum in the designated frequency band (Figure S3B,C). To quantify the relative predominance of 2-Hz over 6-Hz activity, a normalized ratio (2Hz - 6Hz) / (2Hz + 6Hz) was computed for each macrocontact and compared across regions (Figure S3C).

### Mixed-effects models for regional differences in distal phase–amplitude coupling

To test whether distal phase–amplitude coupling to HPC differed between MTL and non-MTL contacts and whether that difference depended on frequency, we pooled distal PAC estimates for the 1-, 2-and 6-Hz bands across macrocontacts and subjects and fitted a mixed-effects model with an MTL×frequency interaction and recording day as a random factor. Coupling values were transformed for normality (Yeo-Johnson) and distance between contacts was included as a covariate:

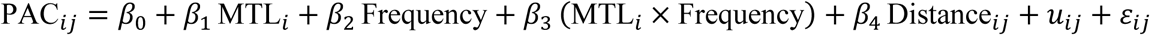

where PAC _𝑖𝑗_ is the transformed distal phase–amplitude coupling value for the pair of distal macrocontact 𝑖 and hippocampal contact 𝑗; MTL_𝑖_ is a binary variable indicating whether contact 𝑖 is in the medial temporal (MTL) or not; 𝐹𝑟𝑒𝑞𝑢𝑒𝑛𝑐𝑦 represents the categorical frequency factor (1-, 2-, or 6-Hz); Distance_𝑖𝑗_ is the Euclidean distance between contact 𝑖 and 𝑗 and controls for inter-contact spacing; 𝑢_𝑖𝑗_ is the random intercept accounting for subject-level variability; and 𝜀_𝑖𝑗_ is the residual error term. From the fitted model we extracted the MTL − non-MTL simple effect at each frequency (estimated difference ± 95% CI) and tested difference-of-differences contrasts (2-Hz versus 1-Hz and 6-Hz) using Wald *t*-tests. This method quantifies whether MTL contacts show selectively greater slow-frequency coupling and whether that regional bias is specific to the 2-Hz band.

### Estimation of phase-gamma pattern synchronization

Phase synchronization between gamma activity recorded from different macrocontacts was quantified from the dispersion of their preferred hippocampal 1-, 2-, and 6-Hz phases. For each hippocampal macrocontact, the mean preferred phase of gamma activity was determined for all extra-hippocampal sites, and the pairwise circular differences between these phases were computed. The average of these pairwise differences was then inverted (2π – Δ phase) so that higher values reflected stronger phase alignment. Synchronization indices were calculated separately for MTL and non-MTL contacts to compare the spatial organization of phase coupling across regions (Figures 3J and S3E,F).

### Detection of hippocampal ripples

Hippocampal ripple events were identified and validated through a multistage procedure combining automated detection, dimensional embedding, and template-based classification.

*Initial ripple detection using overnight sleep recordings:* Hippocampal LFPs recorded during slow-wave sleep were bandpass-filtered between 60 and 180 Hz, and ripple candidates were identified by thresholding the ripple-band amplitude and comparing it to the closest white-matter macrocontact (referred to as control). Candidate events were retained when the amplitude in the ripple band exceeded five standard deviations and was at least twice as strong than in the control macrocontact, and the event lasted for at least one oscillatory cycle. The precise peak times of the filtered signal were extracted to provide a first set of putative ripples for each macrocontact during overnight sleep.

*Dimensional embedding and template generation:* For each recoding night, candidate ripple waveforms were then extracted from the ripple-band–filtered signal (60–180 Hz) using 25-ms snippets centered on the ripple peaks. These band-limited segments, capturing the oscillatory component of each event, were normalized and used as input to Isomap (15 neighbors, intrinsic dimensionality estimated per recording night) to embed the high-dimensional waveforms into a low-dimensional space while preserving their temporal structure ^78^. Events were clustered with k-means, and clusters corresponding to canonical ripple morphologies were manually verified and retained. Then, within each recording night, 200 waveforms from verified clusters were subsampled and averaged to form “super-events”. These “super-events” were pooled across recording nights and clustered again to generate a compact library of representative ripple templates characterized by their average waveform and dominant frequency. Dominant frequency of the templates was estimated from the inverse of the period between two oscillatory peaks in the average ripple-band–filtered signal. Fourteen different templates were retained, ranging from dominant frequency 67.6 Hz to 113.6 Hz.

*Loose detection of putative ripple events*: A permissive detector was then applied to all hippocampal recordings to capture all potential high-frequency bursts in the ripple range (60–180 Hz). The analytic amplitude envelope was smoothed (width = 80 ms, σ = 40 ms) and adaptively thresholded to detect transient events, targeting 20–30 events per minute while constraining event duration (≥20 ms) and separation (≥150 ms). Periods containing IEDs were excluded to prevent contamination by large, transient high-power deflections and to avoid detecting pathological events. Event peaks were realigned using the instantaneous phase of an 80-Hz narrowband signal to ensure consistent phase definition.

*Template matching and white-matter control:* These loosely detected events were then compared to each validated ripple template using cosine similarity. For every event, the highest similarity score and corresponding template were obtained both in the hippocampal contact and in its anatomically defined white-matter control by realigning to the local peak in the ripple-band. Events were classified as genuine ripples only when their match score exceeded 0.85 in the hippocampal macrocontact but remained below this threshold in the control macrocontact. This match score of 0.85 corresponded to the 95^th^ percentile of the distribution of all match scores obtained in white matter macrocontacts, representing a meaningful null distribution for transient, non-specific fast-frequency events. This two-stage validation —requiring both high morphological similarity to verified sleep ripples and spatial specificity relative to the white-matter reference— ensured that the final set of detected events reflected true, locally generated hippocampal ripples.

### Ripple-triggered averages

Ripple-triggered averages were computed to characterize hippocampal LFP, spike train, and spectral dynamics as well as associated gamma activity from the MTL contacts during rest and sleep sessions. For each hippocampal contact, detected events were realigned to the local peak of the ripple-band signal to ensure consistent phase alignment across ripples. Around each realigned peak, ±250 ms of broadband LFP and corresponding time–frequency spectrograms (10–200 Hz, logarithmic spacing) were extracted and averaged across events to obtain mean ripple-locked waveforms (Figures 4A and S4C,G). In Figure 4A, local spike trains and MTL gamma activity were averaged in 24-ms bins, z-scored within contact/neuron, and averaged across ripples (Figure 4A). In Figure S4C, ripples were detected on the macrocontact and realigned to the local peak of the ripple-band signals obtained from the local and the distal tetrodes. In Figure S4G, ripple-triggered averages were computed on the same example hippocampal macrocontact, from events detected in the task rest or during overnight N1 sleep.

### Characterization of ripple central frequency

Hippocampal ripple events were analyzed from LFPs band-pass filtered between 55 and 200 Hz (zero-phase fourth-order Butterworth filter).

Instantaneous phase and amplitude were obtained using the Hilbert transform. Ripple onset and offset were defined as the first and last time points at which the z-scored amplitude exceeded 2. Within these limits, the unwrapped phase was used to compute the total number of oscillatory cycles as the difference between phase values at offset and onset (in degrees) divided by 360. For instance, an unwrapped phase change of 1800° corresponds to five cycles (1800/360 = 5). The ripple central frequency was then calculated by dividing the number of cycles by the event duration in seconds (e.g., 3 cycles / 400 ms = 75 Hz). This analysis was performed separately in rest (Figure S4F,J) and sleep recordings (Figure S4F,H), with sleep events further grouped by sleep stage (wake, N1, SWS, REM).

### Detection of cross-regional neuronal coactivity motifs

Pairwise coactivity motifs were derived from simultaneously recorded single-unit activity during in-burst versus out-of-burst periods (Figures 4D and S4L), task epochs (Figures 4E and S4M–O) and hippocampal ripples. Bursts were detected as described above (see section “*Detection of oscillatory bursts*”) for 1-, 2-and 6-Hz IMFs. For each recording day, units from the MTL were included when at least five well-isolated neurons were available. Spike trains were converted into binned firing-rate matrices using 25-ms bins. We then selected bins overlapping with oscillatory bursts (Figure 4D, viewing and learning sessions concatenated), photograph presentation and its subsequent inter-stimulus intervals in task (viewing versus learning, best versus worst conditions, Figure 4E,F) or ripple events (±200 ms). Control bins were drawn from out-of-burst epochs (Figure 4D) or out-of-ripple intervals of matched duration and distance from ripple events (*data not shown*). Firing rates were z-scored for each neuron, and the influence of overall population activity was removed by regressing out the instantaneous population rate (*P*). Pairwise coactivity between neurons *i* and *j* was defined as the Pearson correlation between their residual binned firing rate, producing one coactivity matrix per condition. To assess the specificity of these patterns, neuron identities were randomly permuted within each time bin, preserving the instantaneous population profile but disrupting pairwise structure. Coactivity matrices computed from the permuted data provided null distributions (Figure S4O). For confirmation, we verified that hippocampal ripples propagated in MTL structures during SWS and N1 sleep (*data not shown*).

### Reactivation of neuronal coactivity motifs in hippocampal ripples

Reactivation was quantified by relating neuronal coactivity motifs expressed during wake (evoked by photographs or during oscillatory bursts) to those measured during pre-and post-learning ripples. For each recording day, the neuron-by-neuron coactivity matrices were vectorized and concatenated across participants to form predictor and response variables (n = 603 coactivity motifs from 6 participants with at least 5 simultaneously recorded MTL neurons). Group-level relationships were estimated using generalized linear models (GLMs).

For each waking condition, the following model (Model 1):

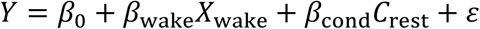

was fitted to quantify the correspondence between waking and rest motifs, where 𝑌 is the vector of ripple coactivity coefficients, 𝑋_wake_ the corresponding waking coefficients (in-burst or out-of-burst, viewing or learning, best or worst recalled associations), and 𝐶_rest_ a categorical factor indicating pre-or post-learning rest. Parameter estimates, 95% confidence intervals, and p-values were obtained from Wald t-tests on fitted GLM coefficients. Model 1 provided β coefficients (𝛽_cond_) and their 95% confidence intervals reported in Figure 4D,F (left panels) and Figure S4L (left panel) and are referred to as “Waking vs rest coactivity (β)”.

Reactivation strength was then defined as the change in the slope linking waking and ripple motifs across rest sessions, estimated from the interaction term in the following model (Model 2):

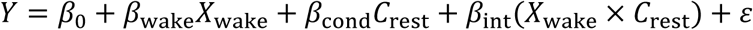

The interaction coefficients (𝛽_int_) and their 95% confidence intervals extracted from Model 2 represent the difference between post-and pre-learning wake–rest slopes and are referred to as “Reactivation (Δ Post – Pre)” in Figure 4D,F (right panels) and Figure S4L (right panel). The *p*-value of these interaction coefficients (𝛽_int_) was used to assess significance of the reactivation in Figure 4D,F (left panels) and Figure S4L (left panel).

Differences in reactivation between waking conditions (in-burst vs out-of-burst, viewing vs learning, best vs worst recalled associations) were tested with a combined model (Model 3) including a three-way interaction term 𝑋_wake_ × 𝐶_rest_ × 𝐶_cond_, where 𝐶_cond_ indexes the waking condition. The *p*-value of this three-way interaction coefficient was used to assess significance in Figure 4E, 4D,F right panels and S4L right panel.

At the subject level (Figure S4M), reactivation was assessed as the partial correlation between waking and ripple coactivity vectors computed separately for each recording day, controlling for the alternate rest session (pre-learning controlling for post-learning rest and vice versa).

### Statistical analysis

Data analyses were conducted using Python version 3.10, incorporating the following packages: DABEST v2023.2.14 ^79^, scikit-learn v1.2.2 and nilearn v0.10.1 ^80^, NumPy v1.24.3^81^, SciPy v1.10.1 ^82^, Stats-Models v0.14.0 ^83^, Matplotlib v3.7.1 ^84^, Pandas v1.5.3 ^85^, and Seaborn v0.11.0 ^86^, MNE-Python v1.5.1 ^73^. Symmetric distribution assumptions underpinned the two-sided statistical tests, visualized using Gardner-Altman and Cumming plots from the DABEST Framework (e.g. Figures 1K, 3J and S2D). These plots illustrate effect sizes by comparing mean or median differences across groups. Each plot consists of two panels: the top (or left) shows raw data distributions with group means ± SEM (unless stated otherwise), and the bottom (or right) shows differences relative to a reference group, calculated from 5,000 bootstrapped samples. Black dots represent the mean (or median), black ticks indicate 95% confidence intervals, and bootstrapped error distribution curves are included. To compare two conditions, bootstrap tests were employed. These tests, which accommodated both paired and unpaired comparisons, estimated the bootstrapped mean difference (either absolute or as a percentage relative to one of the two variables) by resampling the data 100,000 times (unless stated otherwise) with replacement. For paired comparisons, indices were resampled to preserve the relationship between pairs, whereas for unpaired comparisons, each condition was resampled independently. P values for these tests were computed numerically, under the null hypothesis of zero difference. The p value was determined by multiplying the smaller proportion of bootstraps below or above zero by two. Median differences were preferred over mean differences when the original distributions were skewed (visual inspection). In some instances (e.g. Figures 1J, 2D,E, and 3E,F), we visualized the bootstrapped mean (or median) differences using histograms or boxplots, from which the corresponding p values were derived. These histograms do not depict the distribution of raw data points but rather represent empirical estimations of the sampling distribution of the mean difference, obtained by the resampling approach described above. All confidence intervals (95% CI) were calculated via bootstrapping with 100,000 resamples (unless stated otherwise). For each interval, data were resampled randomly with replacement, and the 2.5th and 97.5th percentiles of the bootstrapped distributions determined the lower and upper bounds of the CI. Two-sided *t*-tests or Wilcoxon signed-rank tests were also used to compare conditions, depending on whether normality (assessed by the Shapiro–Wilk test) was met.

Significance of model coefficients was evaluated using two-sided Wald tests.

## Acknowledgements

We thank all subjects and their families for their participation; the staff in the Purpan hospital at the University of Toulouse and at the Pitié-Salpêtrière hospital in Paris for their support; B. Staresina for commenting on a previous version of the manuscript; B. Micklem for technical assistance; all members of the Dupret and Reddy labs for feedback during the project. This work was supported by the Medical Research Council (MRC) UK (programme MC_UU_00003/4 and award MR/W004860/1 to D.D.), the BIAL foundation (award 109/16 to L.R.), and an internal funding from the CerCo (to L.R). A.A.C. is supported by an MRC UK studentship (MC_ST_BNDU_2019). H.C.B. is supported by a UKRI fellowship (MR/W008939/1). The MRC Centre of Research Excellence in Restorative Neural Dynamics is funded by the Medical Research Council UK (award UKRI/MR/B000936/1).

## Author contributions

Conceptualization, A.A.C. and D.D.; Investigation, A.A.C.; Analysis, A.A.C. and D.D.; Methodology, A.A.C., J.C., V.L-d-S., and D.D.; Resources, J.C., R.N-d-S., H.C.B., V.D., M.D., A.D.B., J.C.S., J.A.L., K.L., S.F-V., V.F., V.N., L.V., E.J.B., T.D., L.R., and D.D.; Visualization, A.A.C. and D.D.; Funding acquisition: L.R. and D.D.; Writing – Original Draft, A.A.C. and D.D.; Writing – Reviewing & Editing, A.A.C., J.C., V.L-d-S., R.N-d-S., H.C.B., V.D., M.D., A.D.B., J.C.S., J.A.L., K.L., S.F-V., V.F., V.N., L.V., E.J.B., T.D., L.R., and D.D.; Supervision, D.D.

## Supplementary Figures

**Figure S1.**
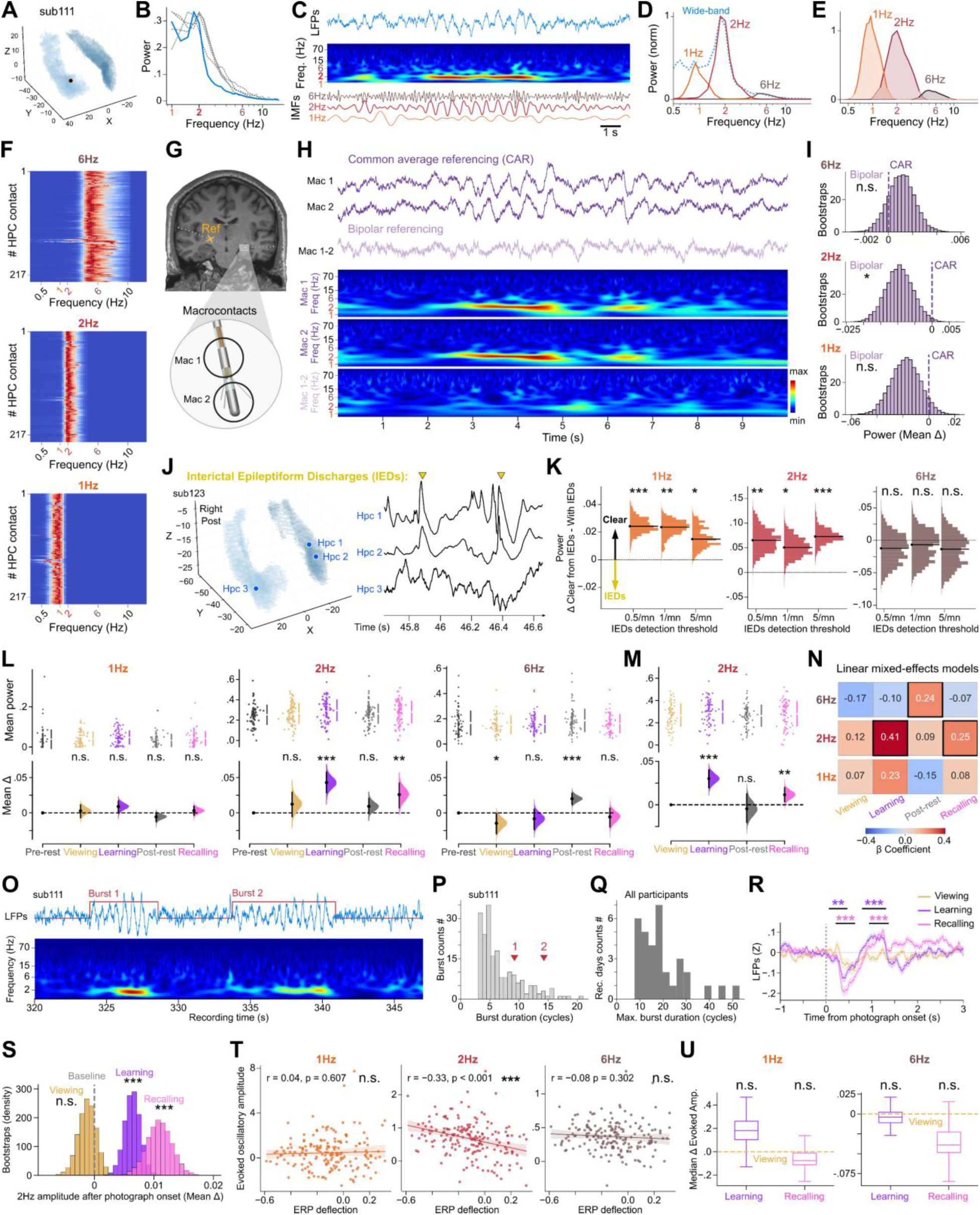
Characterization of slow oscillations in human hippocampus. (A-C) The human hippocampus exhibits 2-Hz oscillations during waking behavior. **(A)** Example 3D projection showing electrode contacts in the hippocampus of one participant. **(B)** Power spectral densities (PSDs) from hippocampal recordings in waking task sessions reveal a prominent oscillation peak around 2-Hz in individual participants (*blue curve*, participant shown in A and C; *gray curves*, other example participants). All contacts included were free of interictal epileptiform discharges (IEDs). **(C)** Example recording showing 2-Hz oscillations (top), corresponding spectrogram (middle), and constituent oscillatory components identified by tailored masked empirical mode decomposition^40^ (intrinsic mode functions, IMFs; see D-F). **(D-F)** Tailored masked empirical mode decomposition of hippocampal LFPs. **(D)** Example PSD of the wide-band signal (dashed line) and corresponding 1-, 2-, and 6-Hz IMFs from a single electrode contact. **(E)** Average PSD across all hippocampal contacts, normalized to maximal 2-Hz power (thick lines: 80% power bands). **(F)** Heatmaps showing power distributions of the 1-Hz (peak [80% power band (PB)]: 1.08 (0.65–1.50) Hz), 2-Hz [peak (80% PB): 2.38 (1.25–3.50) Hz], and 6-Hz [peak (80% PB): 6.15 (3.75–8.50) Hz] IMFs across all hippocampal contacts. **(G-I)** Local referencing reduces detection of slow oscillations. **(G)** T1-weighted MRI showing hippocampal contacts (top) and schematic of local referencing (bottom). Monopolar signals were acquired using a distal white-matter contact and re-referenced either using a common average reference (CAR; median across contacts) or by bipolar subtraction of adjacent contacts (e.g., Mac 1 minus Mac 2). **(H)** Example recording showing LFP traces referenced using CAR (purple) or local bipolar referencing (pink), with corresponding spectrograms. **(I)** Mean difference in hippocampal 1-, 2-, and 6-Hz power after bipolar referencing relative to CAR. **(J,K)** Slow oscillation amplitude decreases at hippocampal contacts with higher interictal discharge rates. **(J)** 3D hippocampal volume showing three recording sites used in **K** (left) and example simultaneous recordings from these contacts (right). Yellow arrowheads: IEDs detected on the Hpc 1 contact. **(K)** Median differences in 1-, 2-, and 6-Hz power between contacts free of IEDs and those with IEDs, plotted as a function of IED-rate threshold. Spearman correlation between IED rate and 2-Hz power: r = - 0.25, P < 0.001; and 6-Hz power: r = 0.06, P > 0.336. **(L-N)** Hippocampal 2-Hz power increases during learning and recall. **(L)** Estimation plot showing mean power differences for hippocampal 1-, 2-, and 6-Hz oscillations across task sessions relative to pre-learning rest. **(M)** Same format as **L**, but relative to viewing. **(N)** Heatmap of β coefficients from a linear mixed-effects model predicting hippocampal 1-, 2-, or 6-Hz power as a function of task session (pre-learning rest as reference), with subject modeled as a random effect; black squares indicate significant coefficients (Wald test, *p* < 0.05). **(O-Q)** Hippocampal slow oscillatory bursts. **(O)** Example raw LFP trace (top) showing 2-Hz oscillatory bursts with corresponding spectrogram (bottom). **(P)** Distribution of burst durations for the contact shown in **O**; arrowheads indicate durations of the two bursts visible in **O**. **(Q)** Distribution of maximal burst duration across participants. **(R-U)** Event-related modulation of hippocampal slow oscillations. **(R)** Group-level average LFPs aligned to photograph onset during viewing, learning, and recall; stronger ERPs are observed during learning and recall. **(S)** Mean differences in post-stimulus 2-Hz amplitude relative to pre-stimulus baseline across task sessions. **(T)** Correlation between ERP deflection and evoked oscillatory amplitudes at 1-, 2-, and 6-Hz during recall. **(U)** Median differences in evoked 1-Hz (left) and 6-Hz (right) amplitude during learning or recall relative to viewing (computed over post-ERP epochs >1 s after photograph onset). Data were analyzed using two-sided paired permutation tests except in **K** and **R** where unpaired tests and cluster-based permutation tests were applied, respectively; ***P < 0.001, **P < 0.01, *P < 0.05; n.s., not significant.

**Figure S2.**
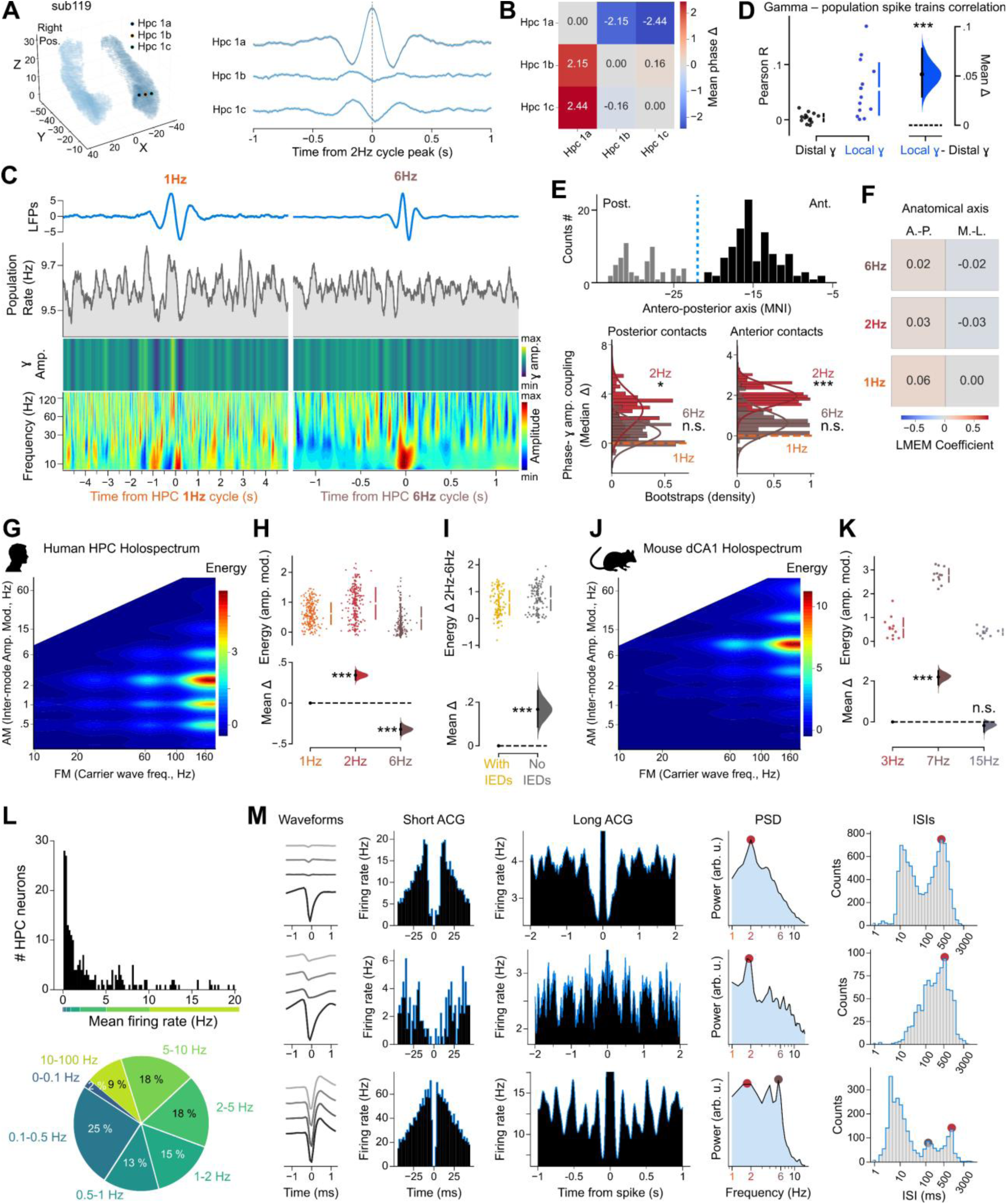
Coupling of neuronal spiking and gamma-band activity to hippocampal slow oscillations. (A,B) Phase reversal of hippocampal 2-Hz oscillations. **(A)** 3D hippocampal volumes showing three linearly arranged recording contacts from the same electrode shaft (left; Hpc 1a, 1b, and 1c) and corresponding average LFPs aligned to hippocampal 2-Hz oscillatory peaks (right), revealing polarity reversal across adjacent contacts. Shaded areas indicate mean ± SEM. **(B)** Heatmap of average phase differences between the three contacts shown in A. The observed reversal across linearly arranged contacts is reminiscent of cross-layer phase shifts described for hippocampal oscillations in animal models^15,39^. **(C-F)** Gamma activity coupling to hippocampal oscillations. **(C)** Average hippocampal LFPs aligned to 1-Hz (left) or 6-Hz (right) phases (blue trace) with corresponding instantaneous population rate, gamma amplitude, and spectrogram (see Figure 2B for comparison). **(D)** Estimation plot showing the difference in instantaneous correlations between neuronal spiking and gamma envelopes recorded at distal versus local macrocontacts. **(E)** Distribution of hippocampal contacts along the antero-posterior axis (top; anterior sites in black, posterior sites in gray) and median phase–amplitude coupling (z-score) differences between 1-Hz and 2-or 6-Hz oscillations across posterior (bottom left) and anterior (bottom right) contacts. Consistent 2-Hz preference is observed in both regions. **(F)** Heatmap of β coefficients from linear regression predicting phase–amplitude coupling (z-score) to 1-, 2-, or 6-Hz oscillations as a function of antero-posterior or medio-lateral contact position. No coefficients reached significance. **(G-K)** Additional Holo-Hilbert spectral analysis further supports gamma amplitude modulation by 2-Hz oscillations in the human hippocampus. **(G)** Holospectrum averaged across time, IMFs, and hippocampal macrocontacts (z-scored), revealing preferential modulation of fast-frequency signals by 2-Hz oscillations in humans. **(H)** Estimation plot showing mean differences in amplitude modulation between 1-Hz and 2-or 6-Hz oscillations across hippocampal macrocontacts. **(I)** Estimation plot showing stronger 2-Hz versus 6-Hz modulation in contacts free of interictal discharges (IEDs) compared with those containing IEDs. **(J)** For comparison, holospectrum averaged over hippocampal CA1 contacts in mice^39^, revealing preferential modulation of fast-frequency signals by ∼7-Hz oscillations. **(K)** Quantification of **J** as in **H**, showing dominant amplitude modulation at ∼7 Hz in mice (n = 12 sessions, 6 mice). **(L,M)** Firing-rate distribution and rhythmicity of hippocampal neurons. **(L)** Mean firing rates of hippocampal neurons follow a log-normal distribution, with most (73%) below 5 Hz. **(M)** Example hippocampal neurons (one per row) exhibiting 2-Hz rhythmicity. From left to right: mean spike waveform across tetrode channels; short-(millisecond) and long-(second) timescale spike autocorrelograms (ACGs); power spectral density (PSD); and inter-spike interval (ISI) distribution. Data were analyzed using two-sided paired permutation tests except in **I** where unpaired tests were applied; ***P < 0.001, *P < 0.05; n.s., not significant.

**Figure S3.**
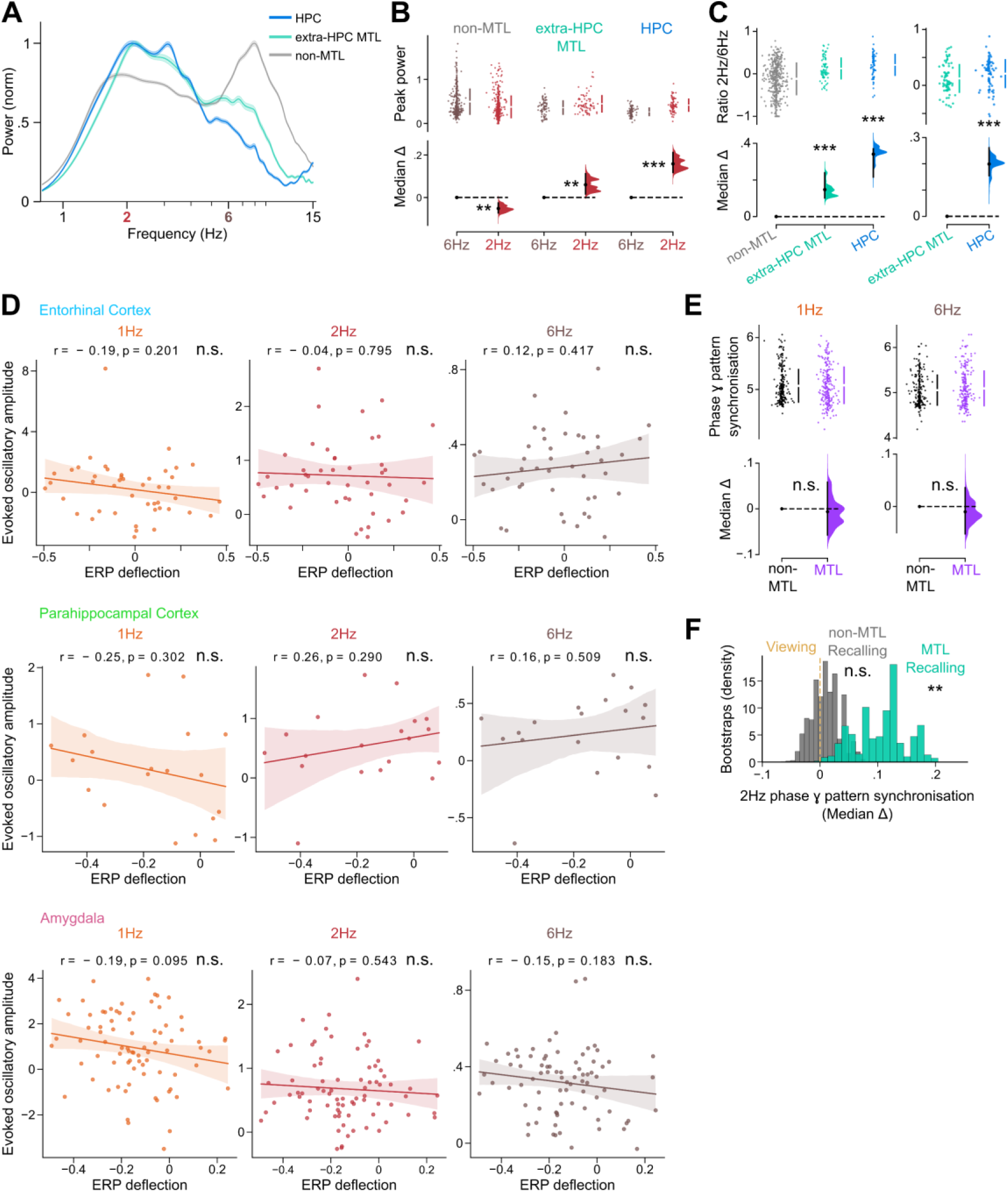
Oscillatory power and gamma-band synchronization across the temporal lobe. (A-C) Slow-frequency oscillations across the temporal lobe. **(A)** Power spectral densities (PSDs) corrected for the aperiodic (1/f) component and averaged across hippocampal (HPC), extra-hippocampal medial temporal lobe (extra-HPC MTL), and non-MTL temporal lobe macrocontacts. All contacts were free of interictal discharges. Shaded areas indicate mean ± SEM. **(B)** Estimation plots showing mean differences between corrected 2-Hz and 6-Hz peak power in the regions shown in **A**. **(C)** Estimation plots showing higher 2-Hz/6-Hz power ratios in hippocampal and extra-hippocampal MTL contacts compared with non-MTL regions (left), and higher 2-/6-Hz power ratios in hippocampal contacts compared with extra-hippocampal MTL contacts (right). **(D)** ERP deflection does not correlate with evoked 2-Hz bursts outside the hippocampus. Correlations between ERP deflection and evoked 1-, 2-, and 6-Hz amplitudes during recall in the entorhinal cortex (top), parahippocampal cortex (middle) and amygdala (bottom). No correlation reached significance. **(E,F)** Cross-regional 2-Hz phase synchronization of gamma-band activity. **(E)** Estimation plot showing median differences in 1-Hz (left) and 6-Hz (right) phase synchronization between MTL and non-MTL gamma-band activity patterns during learning. **(F)** Estimation plot showing median differences in 2-Hz phase synchronization between MTL and non-MTL gamma-band activity patterns during recall relative to viewing. Data were analyzed using two-sided paired permutation tests, except in **C** where unpaired tests were applied; ***P < 0.001, **P < 0.01; n.s., not significant.

**Figure S4.**
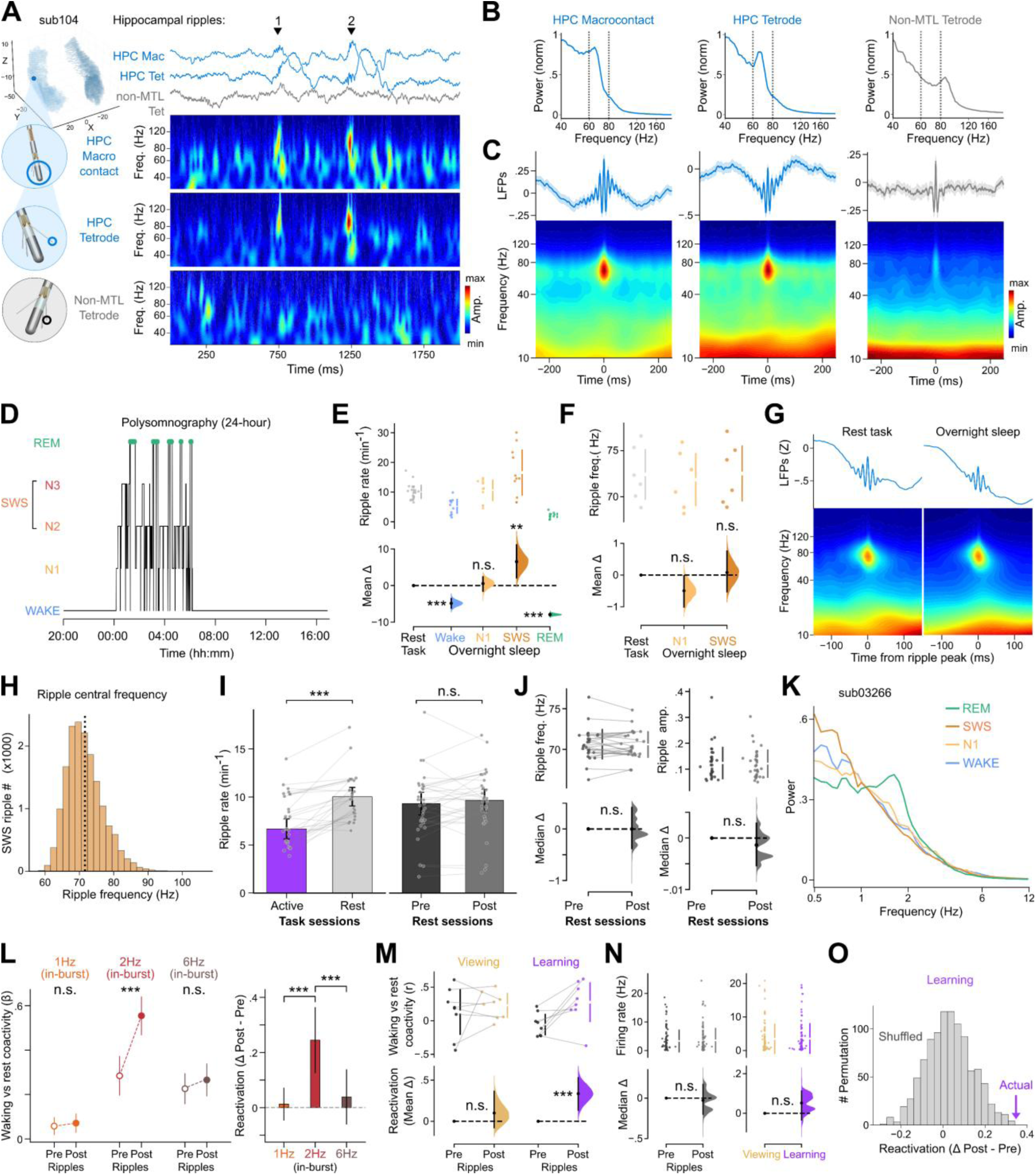
Detection, validation, and control analyses of human hippocampal ripples. (A-C) Ripples detected on hippocampal macrocontacts and tetrodes. **(A)** Example rest recording showing LFP traces and corresponding spectrograms from a hybrid hippocampal electrode (macrocontact and local tetrode) and from a distal, non-MTL tetrode. **(B)** Average PSDs across pre-and post-learning rest sessions showing a peak at ∼70 Hz on the hippocampal macrocontact and tetrode. The distal, non-MTL tetrode shows a peak at ∼85 Hz. **(C)** Ripple-triggered average LFPs and spectrograms aligned to the local peak of the macrocontact or tetrode signals, showing ripples detected on hippocampal macrocontacts and local tetrodes but not on distal, non-MTL tetrodes. **(D-J)** Validation of hippocampal ripples detected during rest task sessions. **(D)** Sleep stages across a 24-h recording session in an example participant. *REM*, rapid eye movement; *SWS*, slow-wave sleep. **(E)** Estimation plot showing mean differences in ripple rate between rest sessions of the memory task and overnight sleep stages. **(F)** Estimation plot showing mean differences in ripple central frequency between rest task sessions and overnight N1 or slow-wave sleep (SWS) sessions, using the same contacts from the same subjects (n = 6 participants). Both the occurrence rate [mean (95% CI): 0.17 (0.15–0.18) Hz] and the central frequency [mean (95% CI): 70.60 (69.92–71.28) Hz] of ripples in rest task sessions were comparable to those recorded during overnight SWS. Ripple occurrence was higher in SWS than during REM sleep and wake, and higher during pre-and post-learning rest than during awake task sessions (viewing, learning, and recall). **(G)** Average LFP traces (top) and spectrograms (bottom) triggered by hippocampal ripples during rest task sessions or overnight sleep (N1), recorded from the same macrocontact in the same participant. **(H)** Distribution of ripple central frequencies across detected SWS ripples, pooled across participants recorded overnight (as in **F**). The detection algorithm identified ripples up to ∼90 Hz; the black vertical dashed line indicates the median frequency (70.9 Hz; mean = 71.6 Hz). **(I)** Bar plots showing higher ripple rates during rest (pre-and post-learning) than during waking task sessions (viewing, learning, recall), with no difference between pre-and post-learning rest. **(J)** Estimation plots showing comparable ripple central frequency (left) and amplitude (right) between pre-and post-learning rest. **(K)** Slow oscillatory activity during REM sleep. Prominent 2-Hz oscillations were observed in the human hippocampus during REM sleep, but not during SWS^63^. Note the elevated <1-Hz power during SWS. **(L-O)** Reactivation of multi-regional MTL coactivity motifs across rhythms and control analyses. **(L)** β coefficients from GLMs quantifying the relationship between in-burst waking events (1-, 2-, and 6-Hz) and ripple coactivity motifs in pre-and post-learning rest (left), and corresponding reactivation strength (right; post-minus pre-learning rest). Significant reactivation was observed only for 2-Hz oscillatory bursts. **(M)** Estimation plots showing matrix-level correlations between viewing-or learning-related coactivity motifs and pre-or post-learning ripples. **(N)** Estimation plots showing similar single-neuron firing rates between pre-and post-learning ripples (left) and between viewing and learning sessions (right). **(O)** Reactivation measured from coactivity motifs computed on actual versus shuffled control spike trains. For each time bin, neuron identities were permuted to disrupt pairwise correlations. Reactivation from actual spike trains exceeded the shuffled distribution. Data were analyzed using two-sided paired permutation tests except in **E**, where unpaired tests were applied, and Wald t-tests on GLM coefficients (**L**, left) or interaction terms (**L** right); ***P < 0.001, **P < 0.01; n.s., not significant.

